# Evaluating ^87^Sr/^86^Sr isotope ratios and Sr mass fractions in otoliths of different European freshwater fish species as fishery management tool in an Alpine foreland with limited geological variability

**DOI:** 10.1101/2021.07.23.453494

**Authors:** Andreas Zitek, Johannes Oehm, Michael Schober, Anastassiya Tchaikovsky, Johanna Irrgeher, Anika Retzmann, Bettina Thalinger, Michael Traugott, Thomas Prohaska

## Abstract

The focus of this study was to assess the potential of otolith microchemistry as a fishery management tool for different European freshwater fish species in an Alpine foreland with a diverse range of different water bodies but low geological variation. ^87^Sr/^86^Sr isotope and Sr/Ca ratios in water samples from 26 habitat sites in a pre-alpine catchment region around lake Chiemsee, Germany, an important region for recreational and economic fisheries, were analysed. ^87^Sr/^86^Sr isotope ratios and the Sr mass fractions in otoliths of 246 fish out of 16 species were determined using (laser ablation) inductively coupled plasma mass spectrometry ((LA)-ICP-MS). Habitats could be discriminated into three distinct strontium isotope regions (SIGs) and seven clusters with characteristic ^87^Sr/^86^Sr isotope and Sr/Ca ratios. The direct comparison of ^87^Sr/^86^Sr isotope ratios in water and otolith samples allowed to identify fish that might have been a) migrating b) transferred from other water bodies or c) stocked from fish farms. Sr/Ca ratios in water and the Sr mass fraction in otoliths were highly correlated, although significant differences between species from the same environment could be documented. Sr mass fractions in sagittae of *Perca fluviatilis* were about 60 % of those in sagittae of *Coregonus spp* and of lapilli of roach *Rutilus rutilus* from the same habitats. Different partition factors for water to otolith Sr/Ca mass fractions were determined for different species. Discrimination of fish otoliths by ^87^Sr/^86^Sr isotope ratios and Sr mass fractions according to habitat clusters was possible with success rates ranging from 92 % to 100 % for cyprinids, European perch *Perca fluviatilis*, whitefish *Coregonus spp*. and European grayling *Thymallus thymallus,* and was 74 % for salmonids. Otolith microchemistry proved to have great potential to serve as a fishery management tool at smaller spatial scales such as in the studied Alpine foreland when considering the limited variation of ^87^Sr/^86^Sr isotope and Sr/Ca ratios, the type and spatial distribution of habitats, and the species and question under investigation.

**Highlights:**

- Otolith microchemistry applied in in area with limited geological variability
- Fish transferred, stocked or migrated were identified
- Regressions between Sr/Ca ratios in water predict Sr mass fractions in otoliths
- Species specific Sr discrimination from water into otoliths
- European freshwater fish species assigned to habitat clusters of origin

## 1. Introduction

The analysis and interpretation of elemental and isotopic fingerprints in fish otoliths has become an important tool for a wide variety of questions related to an improved fisheries management in freshwater systems (Carlson et al., 2017). Questions that have been addressed by this method are e.g. the reconstruction of life histories of fish (Kennedy et al., 2002), the determination of natal origins and movement for designing spatially informed habitat conservation programs and harvest regulations (Carlson et al., 2016), the determination of contributions of different reproduction areas for an improved management and conservation of whole fish communities (Zeigler and Whitledge, 2010; Zeigler and Whitledge, 2011), the management of selected economically relevant fish species (Veinott and Porter, 2005; Walther et al., 2008; Radigan et al., 2018b) and the management of recreational fishery (Veinott et al., 2012) or of endangered fish species (Strohm et al., 2017). Furthermore, elemental and isotopic fingerprints in fish otoliths have been applied to assess the efficiency of ecological rehabilitation measures (Schaffler et al., 2015), the management of invasive species (Blair and Hicks, 2012; Norman and Whitledge, 2015), to identify source and date of introduction of fish from other sources (Munro et al., 2005), to discriminate between hatchery reared and stocked and naturally spawned fish in river systems (Coghlan et al., 2007; Zitek et al., 2010) and study natal homing (Engstedt et al., 2014).

Globally, most freshwater studies applying otolith microchemistry focused on single catchment river systems on a larger scale with significantly varying geology, e.g. to identify natal origins of anadromous salmon species (Kennedy et al., 2000; Barnett-Johnson et al., 2008; Brennan et al., 2015b). Avigliano et al. (2021) tracked migrations of a single freshwater species (*Prochilodus lineatus*) in riverine habitats also on a very large scale in the La Plata basin. Only rarely otolith chemistry was assessed as a potential tool for fishery management for several species on a relatively small scale including different habitat types (Zeigler and Whitledge, 2010; Radigan et al., 2018c).

In Europe otolith microchemistry has been mainly primarily used to investigate diadromous migrations and habitat use of European eel *Anguilla anguilla* (Daverat et al., 2005; Tzeng et al., 2005; Arai et al., 2006), twaite shad *Alosa fallax* (Magath et al., 2013), whitefish *Coregonus maraena* (Gerson et al., 2021) or typical freshwater fish species such as European pike *Esox lucius* which also among those inhabits brackish waters (Rohtla et al., 2012). Studies concerning typical European fish species inhabiting only freshwater habitats are rare (Lenaz et al., 2006; Zitek et al., 2010).

Spatial differences in water chemistry in and among river catchments form the basis for the applicability of this approach, e.g. for allocating fish to specific areas by differences in their otolith chemistry (Elsdon et al., 2008; Zimmerman et al., 2013). Especially the radiogenic ^87^Sr/^86^Sr isotope ratio has been identified as major parameter for tracing fish back to geographic locations (Kennedy et al., 2000; Pouilly et al., 2014), as it is incorporated into living organisms according to its availability without significant fractionation (Graustein, 1989; Capo et al., 1998; Blum et al., 2000). Furthermore, this ratio is independent from other influencing factors that have been linked to changes in the uptake of elements such as temperature, salinity, growth, maturation and genetics (Kennedy et al., 2000). Moreover, the ^87^Sr/^86^Sr isotope ratio is expected to be more or less stable across seasons and years in most river systems (Kennedy et al., 2000; Martin et al., 2013), although also indications of potential interannual variability have been recently indirectly assumed by otolith chemistry of a resident fish species (Brennan et al., 2015a). The term “strontium isotope groups” (SIGs) that represent grouped habitats with differentiable (non-overlapping) ^87^Sr/^86^Sr isotope ratios was coined by Brennan et al. (2015b).

The main cause for the ^87^Sr/^86^Sr isotope ratio variation in freshwater catchments is the underlying geology (Faure and Mensing, 2005). The spatial arrangement of geological units therefore influences the strontium isotopic composition of the water (Brennan et al., 2014). Thus it is possible to differentiate fish from different origin by the ^87^Sr/^86^Sr isotope ratio in their otoliths for e.g. provenancing (Bataille and Bowen, 2012) or fish ecological questions (Kennedy et al., 2000; Barnett-Johnson et al., 2008; Hegg et al., 2013). To a priori determine the potential efficacy of using otoliths to differentiate between fish of different sources, a systematic analysis of the water chemistry differences and building clusters of similar habitats to identify fish according to the geographic variation of water chemistry have been suggested (Wells et al., 2003; Humston and Harbor, 2006).

To optimize the classification of otoliths, ^87^Sr/^86^Sr isotope ratios have often been combined with mass fractions of different elements (Carlson et al., 2017). Especially Sr, Zn, Pb, Mn, Ba and Fe in otoliths have been documented to be consistent with an environmental effect, with Sr/Ca ratio being the most promising parameter in addition to ^87^Sr/^86^Sr isotope ratios for the determination of the geographic origin of fish (Campana, 1999; Avigliano et al., 2021). The Sr/Ca ratio in water has been identified as primary factor influencing otolith Sr/Ca ratio for freshwater and diadromous fish (Brown and Severin, 2009) by the relative uptake of Sr from the water by fish based on its environmental availability and its incorporation in otoliths (Campana, 1999). Farrell and Campana (1996), for example, found that Ca was taken up by 75 % from water and Sr by 88 %. Campana (1999) described a significant relationship between the water and otolith Sr/Ca ratios. Wells et al. (2003) found almost a linear relationship between water Sr/Ca and otolith Sr/Ca ratios in westslope cutthroat trout (*Oncorhynchus clarki lewisi*) collected throughout the Coeur d’Alene watershed in northern Idaho. A very similar relationship was found by Zeigler and Whitledge (2010) for multiple fish species and Strohm et al. (2017) for the potamodromous *Chasmistes liorus* in a cage experiment. The Sr/Ca ratio also has been described to be the most stable element/Ca ratio in habitats over time and the most influential elemental factor related to the discrimination of otoliths (Olley et al., 2011). As Sr substitutes randomly for Ca in otoliths driven by environmental availability (Doubleday et al., 2014), Sr mass fractions in otoliths are conventionally expressed as a ratio of Ca mass fractions as Ca is suggested to vary relatively little within aragonitic otoliths (± 5%) (Secor and Rooker, 2000).

In comparison, Mg/Ca ratios did not show this tendency for a linear relationship between water and otolith (Strohm et al., 2017), most probably because Mg uptake is influenced by other factors (Campana, 1999). In the study of Avigliano et al. (2021) this relationship could not be identified for Ba/Ca ratios neither.

However, species-specific differences in the uptake of Sr from the environment have been documented (Swearer et al., 2003; Friedrich and Halden, 2008). They were interpreted as results of genetic effects (Friedrich and Halden, 2008) leading to physiological differences in regulating the incorporation of elements into otoliths (Swearer et al., 2003). Specifically this is anticipated as an effect of different crystallization processes or the presence of different proteins that would selectively influence the elemental incorporation in otoliths (Melancon et al., 2009). Interspecific variation in otolith microchemistry in fish from same habitats in general is to be expected to be more expressed in “species that are more distantly related, either phylogenetically or ecologically” (Swearer et al., 2003). The Sr/Ca ratio incorporated in otoliths in relation to the environmental availability of Sr and Ca could also be influenced by growth, temperature, and age, which also has been linked back to the change rate of protein synthesis in relation to the crystallization rate of the otolith (Campana, 1999). Partition coefficients between environmental and otolith Sr/Ca ratios are usually used to describe elemental discrimination caused by physiological barriers during elemental uptake and incorporation in otoliths (Campana, 1999). Sr to Ca partition coefficients can be used for predicting otolith Sr/Ca values from water samples and assess the potential influence of different factors on elemental uptake (Stewart et al., 2021).

Moreover, it is relevant which of the three pairs of otoliths in fishes (i.e., asteriscus, lapillus, sagitta) is used for a specific species, as potential differences in their calcium carbonate polymorphs (aragonite, vaterite, calcite) need to be considered (Pracheil et al., 2019). Typically, sagittae and lapilli are made of aragonite, while asterisci are typically made of vaterite (Campana, 1999; Lenaz et al., 2006; Pracheil et al., 2019), and calcite in otoliths in general is much rarer (Campana, 1999). Aragonite otoliths have significantly higher concentrations of Sr as compared to vaterite otoliths which are higher in Mg and Mn (Lenaz et al., 2006; Pracheil et al., 2019).

Considering the above-mentioned factors, the aim of this study was to assess to which degree otolith chemistry is applicable as a fishery management tool to multiple species in an Alpine foreland with a diverse range of different water bodies but an expected low geological variation. The hypotheses were, that (1) differences in the ^87^Sr/^86^Sr isotope and Sr/Ca ratios between the investigated water bodies exist and can be used to determine useful habitat clusters, that (2) species differ regarding the uptake of Sr from the environment, and that (3) otoliths could be assigned to their habitat clusters of origin with a high probability when potential species-specific differences are considered.

## 2. Material and Methods

### 2.1 Study Area

Water and fish samples were taken in Chiemgau, an area located in the Alpine foreland of Bavaria (Germany) in the north of the Tyrolian limestone Alps (Fig. 1). The geological background is a mosaic of fluvial, glacial, hillslope and former lake deposits. In this region 26 different sites situated in 19 different rivers, lakes, ponds and one fish farm within a 50 km range of the biggest lake of Bavaria, lake Chiemsee were sampled for water and fish (Tab. 1 and Tab. A.1). The chosen waterbodies are of high economic importance for freshwater fisheries (e.g. for whitefish*, Coregonus spp*.) or recreational fishery (*Thymallus thymallus*, *Salmo trutta*, *Oncorhynchus mykiss* in rivers and species like *Perca fluviatilis*, *Esox lucius* and *Sander lucioperca* in lake fishery), and also important habitats and spawning grounds for endangered fish species like *Rutilus meidingeri, Hucho hucho* or *Salmo trutta f. lacustris.*

**Figure 1:**
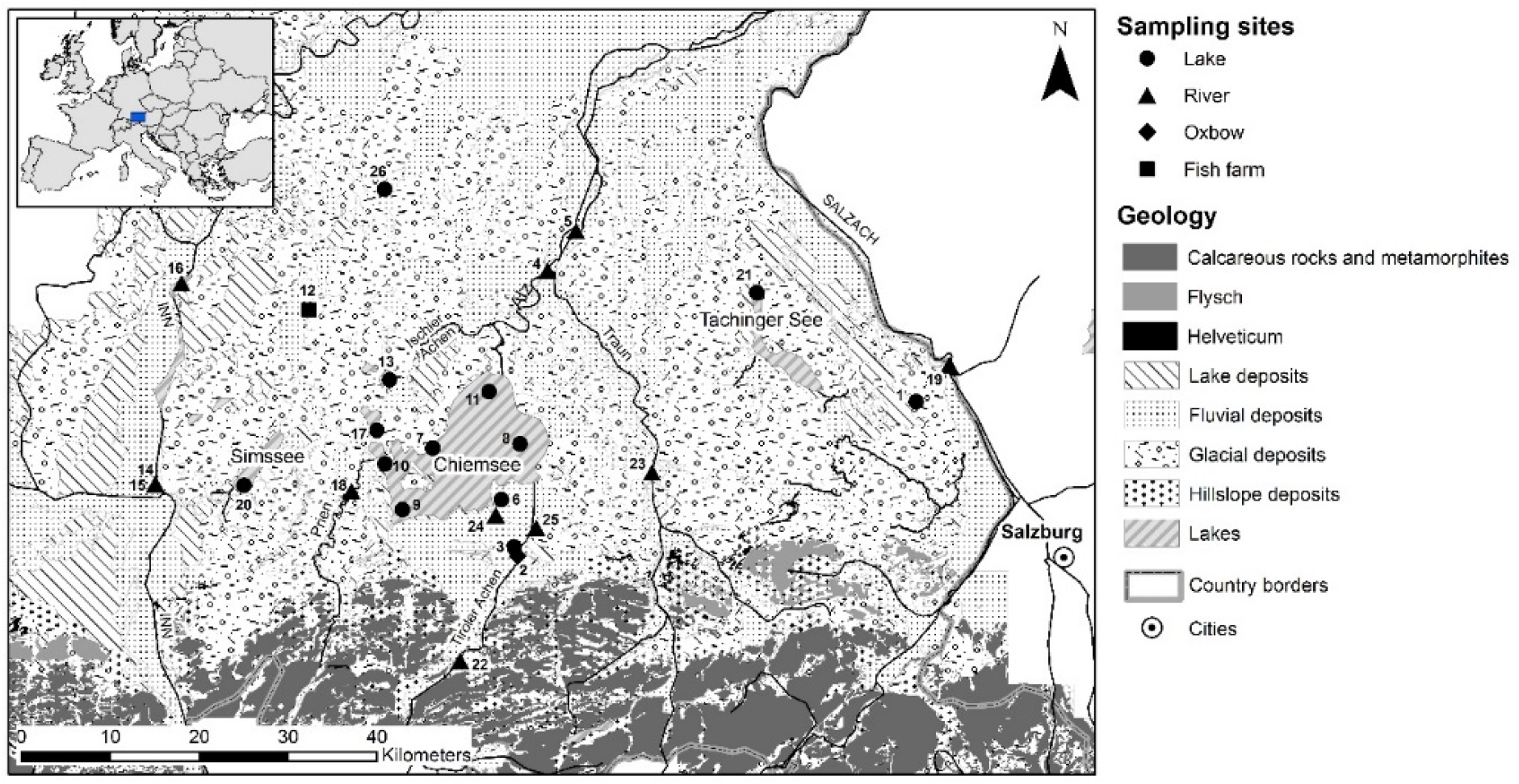
Study area in Southern Bavaria, Germany with its lakes and streams, the simplified underlying geology (data source: Bayerisches Landesamt für Umwelt, www.lfu.bayern.de) and the numbered locations and types of the different sampling points for water and fish; further details can be found in Tab. 1 and Tab A.1

**Table 1:** Sampled water bodies, type and location, expected fish community (*A=Anguillidae*, *Co=Coregonidae, Cy=Cyprinidae, E=Esocidae, G=Gadidae, S=Salmonidae, S(T)=Salmonidae including Thymallus thymallus, P=Percidae*), fish species samples (date of sampling and detailed geographical coordinates can be found in Tab A.1). Most fish samples were sampled directly at the location of the water sample, and if not, they were assigned to the section of the river characterized by the water sample. Lake Chiemsee, Inn and Alz were sampled at multiple locations; all other water bodies were sampled at a single location. For six water bodies (*) the exact location of the fish samples was not available as these fish were captured by fishermen within the investigated lakes, or in one occasion, along the lower course of the River Prien; further details can be found in Tab A.1).

| No | Water Body | Type | Location | Community | Number of individuals per fish species analyzed (n) |
| --- | --- | --- | --- | --- | --- |
| 1 | Abtsdorfer See | Lake | Laufen | Cy, E, S, P | *Rutilus rutilus* (4), *Perca fluviatilis* (4) |
| 2 | Altwasser Osterbuchberg | Oxbow | Grabenstätt | Cy, E, S, P | *Tinca tinca* (1), *Perca fluviatilis* (3) |
| 3 | Almfischerweiher | Pond | Übersee | Cy, E, S, P | *Perca fluviatilis* (7) |
| 4 | Alz upstream Traun entry | River | Altenmarkt | A, Cy, E, S, P | *Barbus barbus* (2), *Esox lucius* (3), *Perca fluviatilis* (5), *Anguilla anguilla* (1) |
| 5 | Alz downstream Traun entry | River | Trostberg (downstr. Altenmarkt) | A, Cy, E, S, P | *Perca fluviatilis* (4) |
| 6 | Baggerweiher Übersee | Pond | Übersee | Cy, E, S, P | *Scardinius erythrophthalmus* (5), *Rutilus rutilus* (8), *Abramis brama* (1) |
| 7 | Chiemsee | Lake | Chieming | A, Co, Cy, E, G, S, P | *Rutilus rutilus* (4), *Coregonus spp* (5), *Perca fluviatilis* (7) |
| 8 | Chiemsee | Lake | Felden | A, Co, Cy, E, G, S, P | *Abramis brama* (3) |
| 9 | Chiemsee | Lake | Fraueninsel | A, Co, Cy, E, G, S, P | *Rutilus rutilus* (4), *Perca fluviatilis* (4), *Coregonus spp* (5), *Lota lota* (1) |
| 10 | Chiemsee | Lake | Prien | A, Co, Cy, E, G, S, P | *Rutilus rutilus* (10), *Perca fluviatilis* (9), *Coregonus spp* (7) |
| 11 | Chiemsee | Lake | Seebruck | A, Co, Cy, E, G, S, P | *Rutilus rutilus* (5), *Perca fluviatilis* (5), *Coregonus spp* (5), *Salmo trutta* (1) |
| 12 | Fischzucht Kreissig | Fish Farm | Amerang | Cy, S | *Salvelinus umbla* (4) |
| 13 | Hartsee* | Lake | Eggstätt | A, Co, C, E, S, P | *Rutilus rutilus* (3), *Perca fluviatilis* (2), *Scardinius erythrophthalmus* (1) |
| 14 | Inn Rosenheim upstr. Mangfall entry | River | Nussdorf | Cy, E, G, S(T), P | *Thymallus thymallus* (1), *Oncorhynchus mykiss* (2) |
| 15 | Inn Rosenheim upstr. Mangfall entry | River | Neubeuern | Cy, E, G, S(T), P | *Thymallus thymallus* (5) |
| 16 | Inn upstream Wasserburg | River | Griesstätt | Cy, E, G, S(T), P | *Scardinius erythrophthalmus* (3), *Thymallus thymallus* (1) |
| 17 | Langenbürgner See* | Lake | Breitbrunn | A, Co, C, E, S, P | *Rutilus rutilus* (2), *Perca fluviatilis* (5), *Scardinius erythrophthalmus* (1) |
| 18 | Prien* | River | Prien | Cy, Sa | *Salmo trutta* (1) |
| 19 | Salzach | River | Laufen | Cy, E, G, S(T), P | *Thymallus thymallus* (2), *Lota lota* (5) |
| 20 | Simssee* | Lake | Riedering | A, Co, C, E, S, P | *Perca fluviatilis* (7), *Coregonus spp* (5), *Rutilus rutilus* (3), *Scardinius erythrophthalmus* (1) |
| 21 | Tachinger See* | Lake | Taching | A, Co, C, E, S, P | *Perca fluviatilis* (4), *Scardinius erythrophthalmus* (8) |
| 23 | Traun | River | Altenmarkt | Cy, E, G, S, P | *Salmo trutta* (1) |
| 24 | Überseer Bach | River | Übersee | A, Cy, E, S, P | *Salmo trutta* (4), *Esox lucius* (3), *Leuciscus leuciscus* (4) |
| 25 | Weißbach | River | Grabenstätt | Cy, S | *Salmo trutta* (4) |
| 26 | Weitsee* | Lake | Schnaitsee | Cy, E, P | *Scardinius erythrophthalmus* (3), *Rutilus rutilus* (1), *Perca fluviatilis* (1) |

**Table 2:** Lengths (minimum, maximum, mean, standard deviation - SD) and total number of individuals from different fish species sampled for otoliths from different water bodies.

| Species | Family | Min. (cm) | Max (cm) | Mean (cm) | SD | n |
| --- | --- | --- | --- | --- | --- | --- |
| *Abramis brama* (Ab) | Cyprinids | 17.0 | 17.0 | 17.0 | - | 4 |
| *Anguilla anguilla* (Aa) | Anguillids | 53.0 | 66.0 | 58.8 | 5.1 | 6 |
| *Barbus barbus* (Bb) | Cyprinids | 35.5 | 36.0 | 35.8 | 0.4 | 2 |
| *Coregonus spp.* (Cspp) | Coregonids | 23.8 | 37.0 | 29.7 | 3.7 | 39 |
| *Esox lucius* (El) | Esoxids | 19.0 | 36.2 | 25.8 | 6.8 | 6 |
| *Leuciscus leuciscus* (Lle) | Cyprinids | 14.0 | 21.1 | 17.6 | 3.8 | 4 |
| *Lota lota* (Ll) | Gadids | 13.2 | 46.5 | 29.1 | 12.4 | 9 |
| *Oncorhynchus mykiss* (Om) | Salmonids | 15.1 | 34.0 | 23.0 | 10.0 | 6 |
| *Perca fluviatilis* (Pf) | Percids | 6.4 | 24.0 | 14.4 | 5.0 | 71 |
| *Rutilus rutilus* (Rr) | Cyprinidae | 7.8 | 30.0 | 19.1 | 7.2 | 44 |
| *Salmo trutta* (St) | Salmonids | 11.5 | 33.5 | 22.8 | 8.0 | 11 |
| *Salvelinus umbla* (Su) | Salmonids | 29.0 | 32.0 | 30.5 | 1.3 | 4 |
| *Scardinius erythrophthalmus* (Se) | Cyprinids | 7.6 | 16.1 | 12.3 | 2.5 | 27 |
| *Squalius cephalus* (Sc) | Cyprinids | 6.8 | 18.0 | 11.0 | 6.1 | 3 |
| *Thymallus thymallus* (Tt) | Salmonids | 28.5 | 52.0 | 41.8 | 9.6 | 9 |
| *Tinca tinca* (Tti) | Cyprinids | 26.0 | 26.0 | 26.0 | - | 1 |
| Total |  | 6.4 | 66.0 | 21.9 | 12.1 | 246 |

### 2.2 Water Sampling

Between March 2011 and October 2014 water samples were taken from the described 26 sites from 19 different water bodies. Most of the water bodies were sampled at a single location while at the rivers Inn, Tiroler Ache and Alz in different sections, up- and downstream of important tributaries, water samples were taken. Due to its size, Lake Chiemsee was sampled at seven different locations (Tab. 1 and Tab A.1).

At each sampling site, 100 mL triplicates of water samples were collected in acid-washed polyethylene bottles (pre-cleaned with HNO_3_ (*w* = 10 %) and HNO_3_ (*w* = 1 %) and rinsed with deionized water) using a hand-dipping device to which the open bottles could be mounted. This device allowed sampling from the shore without entering the water from steeper shoreline slopes, also excluding the risk of contamination by material mobilized from the ground. Each sample was collected approximately 3 m off-shore and about 10-20 cm below the water surface. At Lake Chiemsee at Chieming, Felden, Prien and Seebruck water samples were taken at 1 m and 12 m depth using a Schindler-Patalas water sampler (Uwitec, Tiefgraben, Austria) from which three subsamples were bottled. Water samples were kept in a cool box during transport and stored in the laboratory at −28 °C until further treatment.

### 2.3 Fish Sampling

Fish were sampled between August 2013 and April 2014 from 26 sampling sites in the vicinity of the associated water sampling site by fishermen (Tab. 1 and Tab. A.1). At Lake Chiemsee and at the rivers Alz and Inn fish were collected from multiple different locations, while in all other waters the fish samples originated mostly from one site. The fish were caught either by electric fishing, gill nets or fishing-rods. For some fish samples, only heads or carcasses (head with spine and caudal fin without fillets and entrails) were available from fishermen, so lengths were either provided by the fishermen or estimated by otolith growth functions. A total of 246 individuals out of 16 species (salmonid species such as *Oncorhynchus mykiss, Salvelinus umbla* and *Salmo trutta f. fario* and *Thymallus thymallus*, and one lake trout *Salmo trutta f. lacustris,* cyprinid species such as *Abramis brama*, *Barbus barbus, Leuciscus leuciscus, Squalius cephalus, Rutilus rutilus, Scardinius erythrophthalmus* and *Tinca tinca*, percids such as *Perca fluviatilis*, the esocid species *Esox lucius*, the coregonid species *Coregonus spp.*, the gadid species *Lota lota and the* anguillid species *Anguilla anguilla* that have been stocked in the past in several lakes) were included for analysis. Species being sampled were the most abundant species in the region. Fish were kept at −28 °C before otolith extraction.

### 2.4 Otolith preparation and analysis

To remove the otoliths, fish were defrosted and total length (cm) and weight (g) were measured, except for samples where only heads were available. Fish heads were split using chirurgical scissors and the lapilli (cyprinids) and sagittae (non-cyprinids) were removed using metal forceps. Otoliths were cleaned from adherent tissue in a petri dish filled with reagent grade I water (18.2 MΩ cm, TKA-GenPure ‘Part of Thermo Fischer Scientific’, Niederelbert, Germany), transferred into polypropylene vials containing reagent grade I water, sonicated for 5 minutes in an ultrasonic bath (Model No. UD80SH-2L, Eumax Technology Ltd., Shenzhen, China) and air-dried on black clean room towels. Microscope images of all otoliths were taken sulcus side up- and down (Sagittae) and from ventral and dorsal sides of the Lapilli using a binocular microscope (S63T Trinocular Pod (8-50x), China) connected to a digital camera (ProgRes^®^ CT3, Jenooptik, Germany). Subsequently otoliths were sonicated again for 3 minutes, air dried and stored in pre-cleaned polypropylene vials until further processing.

To prepare otoliths for the laser ablation inductively coupled plasma mass spectrometry (LA-ICP-MS) analysis, they were transferred to individually labeled cut pieces of microscope slides (12 mm x 12 mm) and affixed with thermoplastic glue (Crystalbond™ 509, Aremco Products Inc., New York, USA) heated on a heating plate (Type RCT basic, IKA Labortechnik, Staufen, Germany) with sulcus side up. After this step the mounted otoliths were transferred to larger glass plates (95 mm x 113 mm) in groups of 60 for their analysis by LA-ICP-MS. The size of the glass plate was hereby chosen to fit into the laser ablation chamber, and the otoliths were placed in about 1 cm distance from the glass side edge due to the restricted operational movement of the laser. Another proportion of the glass plate was left empty for attaching the standard reference materials directly before measurement. The plates were stored in polyethylene zip bags until further analysis. Measurements of elemental content and ^87^Sr/^86^Sr isotope ratios were performed consecutively in parallel on small flat areas of the upper side of the otoliths for a length of 150 μm for elemental content and 450 μm for ^87^Sr/^86^Sr isotope ratios using a laser ablation system (NWR193, Electro Scientific Industries, Portland, USA) coupled to a quadrupole ICP-MS (NexION 350D, PerkinElmer, Waltham, USA) for multi-elemental analysis followed by a coupling to a multicollector inductively coupled plasma mass spectrometer (MC ICP-MS, NU Plasma HR, Nu Instruments Ltd., Wrexham, UK) for ^87^Sr/^86^Sr isotope ratio analysis. Before the multi-elemental analysis, a pre-ablation (spot size = 75 μm, scan speed = 10 μm s^−1^, repetition rate = 10 Hz and energy set to 1 %) was conducted to remove potential contaminations. However, this step was skipped later for the ^87^Sr/^86^Sr isotope ratio analysis, as no differences in the data of subsequent line scans could be found. Laser parameters for the analysis of ^88^Sr and ^43^Ca using the quadrupole ICP-MS were 50 μm spot size, 5 μm s^−1^ scan speed, 10 Hz repetition rate and 50 % energy. For ^87^Sr/^86^Sr isotope measurements using the MC ICP-MS a spot size of 100 μm and a repetition rate of 15 Hz were used. Laser energy typically ranged between 20-30 % and was adopted to achieve a minimum signal intensity of 1 V with maximum intensities of about 9 V for ^88^Sr on the MC ICP-MS. A membrane desolvating system (DSN-100, Nu Instruments Ltd., Wrexham, UK) was interconnected between the laser to the MC ICP-MS prior to injection to allow the continuous introduction of HNO_3_ (*w* = 2 %) solution into the output of the laser ablation cell and a fast switch between laser ablation and solution-based analysis. In-house pressed pellets of the certified reference materials (CRM) bone ash NIST SRM 1400 (National Institute of Standards and Technology, Gaithersburg, MD, USA) and fish otolith powder FEBS-1 (National Research Council of Canada, Ottawa, Canada) were used for elemental quantification and ^87^Sr/^86^Sr isotope ratio method evaluation respectively. The CRM are certified for Ca and Sr content whereas values for ^87^Sr/^86^Sr isotope ratios have been determined in several studies for FEBS-1 (Yang et al., 2011; Zimmermann et al., 2019) and NIST SRM 1400 (Romaniello et al., 2015; Zimmermann et al., 2019). Data reduction was done according to established procedures (Irrgeher et al., 2016; Prohaska et al., 2016; Retzmann et al., 2019).

### 2.5 Water sample analysis

All water samples were defrosted, filtered via a cellulose acetate filter membrane (Minisart 0.45 μm syringe filter units, Minisart, Sartorius, Göttingen, Germany) and acidified to *w* = 2 % HNO_3_ and diluted as required prior to the elemental measurements. The Sr and Ca elemental mass fractions were measured in standard mode using a quadrupole ICP-MS (ELAN DRC-e or NexION300D, Perkin Elmer, Waltham, MA, USA). For the quantification of the elemental mass fractions of Sr and Ca external calibration standards (multielemental standard, Multi VI, Merck, Darmstadt, Germany) were used. Obtained elemental signals were blank corrected, normalized to indium used as internal normalization standard (single element ICP-MS standard, CertiPur, Merck, Darmstadt, Germany) and quantified via an external multi-point calibration. Results were validated using in-house quality control standards. SLRS-5 river water certified reference material (NRC, Canada) was used for method validation along with all sample preparation steps. Elemental ratios are elemental amount ratios.

Prior to ^87^Sr/^86^Sr isotope ratio measurements, Sr/matrix separation using a Sr-specific extraction resin (Sr Spec Resin, Triskem, Bruz, France) was performed to remove the isobaric interferences of ^87^Rb on ^87^Sr (Horsky et al., 2016) as well as Ca-based polyatomic interferences that form in the plasma and potentially occur on all nominal Sr isotope masses (Zimmermann et al., 2019). The ^87^Sr/^86^Sr isotope ratios of the water samples were measured with a MC ICP-MS instrument (Nu Plasma HR, Nu Instruments, Wrexham, UK). The ^87^Sr/^86^Sr isotope ratios were corrected for instrumental isotopic fractionation using external intra-elemental calibration against the NIST SRM 987 in a standard-sample bracketing mode (Irrgeher and Prohaska, 2015; Horsky et al., 2016). The ^87^Sr/^86^Sr isotope ratios in this study are *n*(^87^Sr)/*n*(^86^Sr) isotope-amount ratios, which is the correct notation according to IUPAC guidelines (Coplen, 2011). Uncertainties were determined using the Kragten approach according to EUROCHEM (Ellison and Williams, 2012; Horsky et al., 2016). All uncertainties in this work correspond to the expanded uncertainties (*U*, *k* = 2) corresponding to a coverage factor of 95 %. In accordance to EURACHEM guidelines, measurement values have to be considered equal when they overlap within limits of uncertainty (Ellison and Williams, 2012). All instruments were optimized for maximum sensitivity and signal stability.

### 2.6 Data analysis

To identify habitat clusters with similar ^87^Sr/^86^Sr isotope and Sr/Ca mass fraction ratios in water samples a three-step approach was taken. Firstly strontium isotope groups (SIGs) after Brennan et al. (2015b) were built considering the expanded uncertainties (*U*, *k*=2). Hereby habitat groups were identified that could be distinguished only by their ^87^Sr/^86^Sr isotope ratios. SIG creation was optimized towards a reduction of transition habitats. Secondly, according to Wells et al. (2003) and Zitek et al. (2010), clusters of similar habitat characteristics were explored using the Ward and Nearest Neighbour algorithms (IBM SPSS Statistics for Windows, Version 24.0) with values standardized to 1 to balance the different dimensions of the two variables. In a third step cluster results were inspected and refined following an approach suggested by Kern et al. (2017) based on analytical considerations related to the defined SIGs in relation to the Sr/Ca values. Finally, the resulting habitat clusters were then interpreted based on the types of waterbodies they included.

To identify fish that might have been stocked, transferred between water bodies or migrated from a nearby river stretch we followed a similar approach as Kennedy et al. (2000) to identify fish that have been moved, so called “movers”. For this purpose ^87^Sr/^86^Sr isotope ratio of all 246 otoliths were plotted in relation to ^87^Sr/^86^Sr isotope ratios of water samples. When the Sr isotope ratio in otoliths was overlapping within the expanded uncertainties *U* (*k*=2) (according to Ellison and Williams (2012)) with that of the water of origin they were considered as originating from the associated habitat. In case, the ^87^Sr/^86^Sr isotope ratio of otoliths and water of origin was different, additional information was collected for interpretation. If the original source could not be retrospectively determined, they were excluded from subsequent analysis of the relation between Sr/Ca in water and the Sr elemental mass fractions in otoliths.

In a next step, the relation between water Sr/Ca and the Sr elemental mass fractions in the reduced set of 226 otoliths was first assessed for each species individually and secondly for all cyprinids and salmonids combined. For this purpose, Spearman rank correlations between Sr/Ca ratio in water and the Sr elemental mass fraction in otoliths were performed and linear regressions were built to display the functional relationship of the Sr/Ca ratio in water to the Sr mass fractions in otoliths. Species-specific differences of the Sr elemental mass fraction in otoliths from fish inhabiting the same habitats were assessed by comparative analysis using the Kruskal-Wallis test followed by Dunn’s post-hoc test without Bonferroni correction. Significance levels were set to 0.05, with *P*-values between 0.05 and 0.1 being noted as showing a strong tendency towards significance. Results of the tests were displayed together with box plots of the Sr elemental mass fractions in otoliths of the different species. These statistical analyses were conducted using IBM SPSS Statistics for Windows, Version 24.0 (Released 2016, Armonk, NY: IBM Corp.).

To further describe the species specific differences in Sr uptake from water of fish living in the same environment, partition coefficients according to Campana (1999) were calculated per species or family with elemental discrimination D_Sr:Ca_ = (Sr:Ca)_otolith_/(Sr:Ca)_water_. Otolith Sr mass fractions in otoliths were converted to Sr/Ca mass fraction ratios (μg g^−1^) using the stoichiometric mass fraction of Ca in aragonite (400.436 μg Ca g^−1^ CaCO_3_) (Whitledge et al., 2019).

To assess the potential to discriminate otoliths according to the determined habitat clusters, first cluster membership numbers were assigned to the respective otolith samples. As recommended by Gao et al. (2013) for otolith isotope data, a non-parametric discriminant analysis as using the DISCRIM procedure of SAS (2007) (kernel = Epanechnikov, radius = 1) was applied to test to which degree the otoliths could be discriminated according to their habitat clusters using standardized values of the ^87^Sr/^86^Sr isotope and Sr mass fractions. Analyses were run for *P. fluviatilis, Coegonus. spp, T. thymallus* separately and for all cyprinids and all salmonids (except *T. thymallus*) combined. The total percentage of correctly classified otoliths was determined, and the ^87^Sr/^86^Sr isotope ratios and the Sr/Ca mass fraction ratios of the water bodies and the result of the otolith discriminant analyses were plotted side by side along with their cluster membership including the associated uncertainties and incorrect cluster assignments. For each species or family group only those habitats and clusters were considered and displayed where an occurrence could be expected.

## 3. Results

### 3.1 Determination of habitat clusters

In a first step three clearly distinguishable habitat strontium isotope groups (SIGs) were identified based on the ^87^Sr/^86^Sr isotope ratios (Fig. A.1). The ^87^Sr/^86^Sr values in these SIGs (considering the expanded uncertainties) ranged from 0.7076-07083 (SIG 1, n=4 water bodies), 0.7083-0.7091 (SIG 2, n=13 water bodies) and 0.7092-0.7100 (SIG 3, n= 3 water bodies). Seven water bodies could not be unambiguously assigned to a SIG because of overlapping uncertainties. Fig. A.2 shows the combination of the determined SIGs with the Sr/Ca values of water samples.

Based on ^87^Sr/^86^Sr isotope and Sr/Ca ratios in water samples from all 26 habitats where also fish were sampled plus one additional habitat added later (mouth of river Inn, No. 27) could be grouped into 7 habitat clusters (Fig. 2).

**Figure 2:**
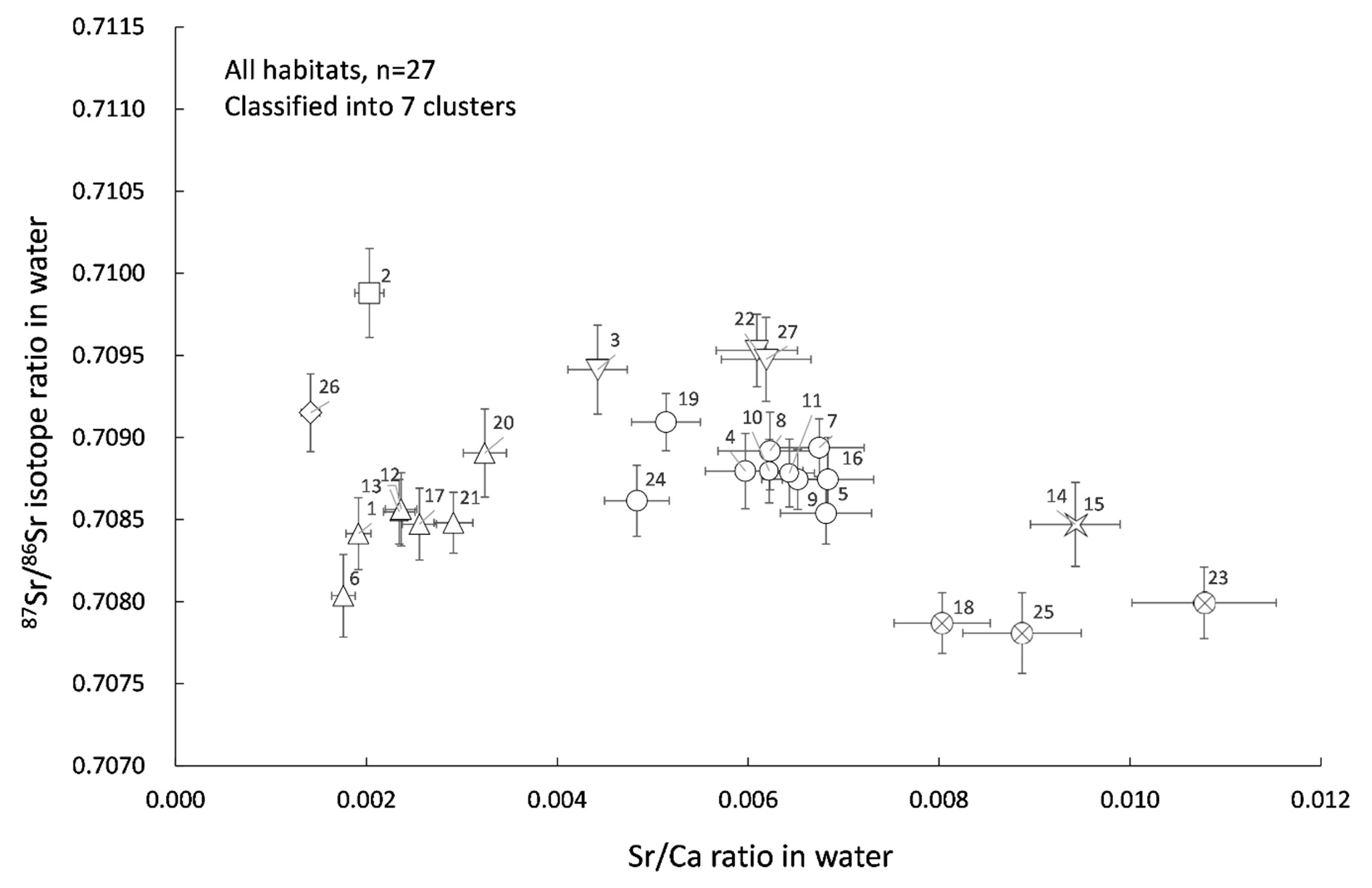
Habitat clusters (n=7) of all 26 sampled habitats plus one habitat added later (mouth of River Inn, No. 27 (Kendlbacher, 2013)) based on SIGs, Ward clustering and the subsequent refinement; triangle = cluster 1, circle = cluster 2, diamond = cluster 3, square = cluster 4, flipped triangle = cluster 5, star = cluster 6, circle with cross = cluster 7.

Cluster No. 1 contained only lacustrine habitats surrounding Lake Chiemsee together with a fish farm but no Lake Chiemsee habitats, cluster No. 2 contained all Chiemsee habitats together with a variety of river habitats, cluster No. 6 and No. 7 only contained only riverine habitats (Tab. 3). Some of the clusters (No. 3 and No. 4) only represented single lacustrine habitats with distinct ^87^Sr/^86^Sr isotope and Sr/Ca ratios.

**Table 3:** Habitat clusters with their associated fish communities (*A=Anguillidae*, *Co=Coregonidae, Cy=Cyprinidae, E=Esocidae, G=Gadidae, S=Salmonidae, S(T)=Salmonidae including Thymallus thymallus, P=Percidae*).

| Cluster number (symbol) | Water body (habitat number) | Type | Fish community |
| --- | --- | --- | --- |
| 1 (triangle) | Abtsdorfer See (1) | Lake | Cy, E, S, P |
|  | Baggerweiher Übersee (6) | Pond | Cy, E, S, P |
|  | Fish farm Kreissig (12) | Fish farm | Cy, Sa |
|  | Hartsee (13) | Lake | A, Co, C, E, S, P |
|  | Langenbürgner See (17) | Lake | A, Co, C, E, S, P |
|  | Simssee (20) | Lake | A, Co, C, E, S, P |
|  | Tachinger See (21) | Lake | A, Co, C, E, S, P |
| 2 (circle) | Alz upstream Traun entry (4) | River | A, Cy, E, S, P |
|  | Alz downstream Traun entry (5) | River | A, Cy, E, S, P |
|  | Chiemsee (7) | Lake | A, Co, Cy, E, G, S, P |
|  | Chiemsee (8) | Lake | A, Co, Cy, E, G, S, P |
|  | Chiemsee (9) | Lake | A, Co, Cy, E, G, S, P |
|  | Chiemsee (10) | Lake | A, Co, Cy, E, G, S, P |
|  | Chiemsee (11) | Lake | A, Co, Cy, E, G, Sa, P |
|  | Inn upstream Wasserburg (16) | River | Cy, E, G, S(T), P |
|  | Salzach (19) | River | Cy, E, G, S(T), P |
|  | Überseer Bach (24) | River | A, Cy, E, S, P |
| 3 (diamond) | Weitsee (26) | Lake | Cy, E, P |
| 4 (square) | Altwasser Osterbuchberg (2) | Oxbow | Cy, E, S, P |
| 5 (flipped triangle) | Almfischerweiher (3) | Pond | Cy, E, S, P |
|  | Tiroler Achen (22) | River | Cy, E, G, S, P |
|  | Mouth of River Inn (27) | River | Cy, E, G, S(T), P |
| 6 (star) | Inn Rosenheim (14) | River | Cy, E, G, S(T), P |
|  | Inn Rosenheim (15) | River | Cy, E, G, S(T), P |
| 7 (circle with cross) | Prien (18) | River | Cy, S |
|  | Traun (23) | River | Cy, E, G, S, P |
|  | Weißbach (25) | River | Cy, S |

### 3.2 ^87^Sr/^86^Sr isotope ratios in otoliths

Otoliths from 26 fish sampling sites showed ^87^Sr/^86^Sr isotope ratios between 0.70724 (± 0.00042 *U*, *k*=2) and 0.71440 (± 0.00042 *U*, *k*=2), and were matched with ^87^Sr/^86^Sr isotope ratios of water ranging from 0.70781 (± 0.00025 *U*, *k*=2) to 0.70988 (± 0.00025 *U*, *k*=2) (Fig. 3).

**Figure 3 a-c:**
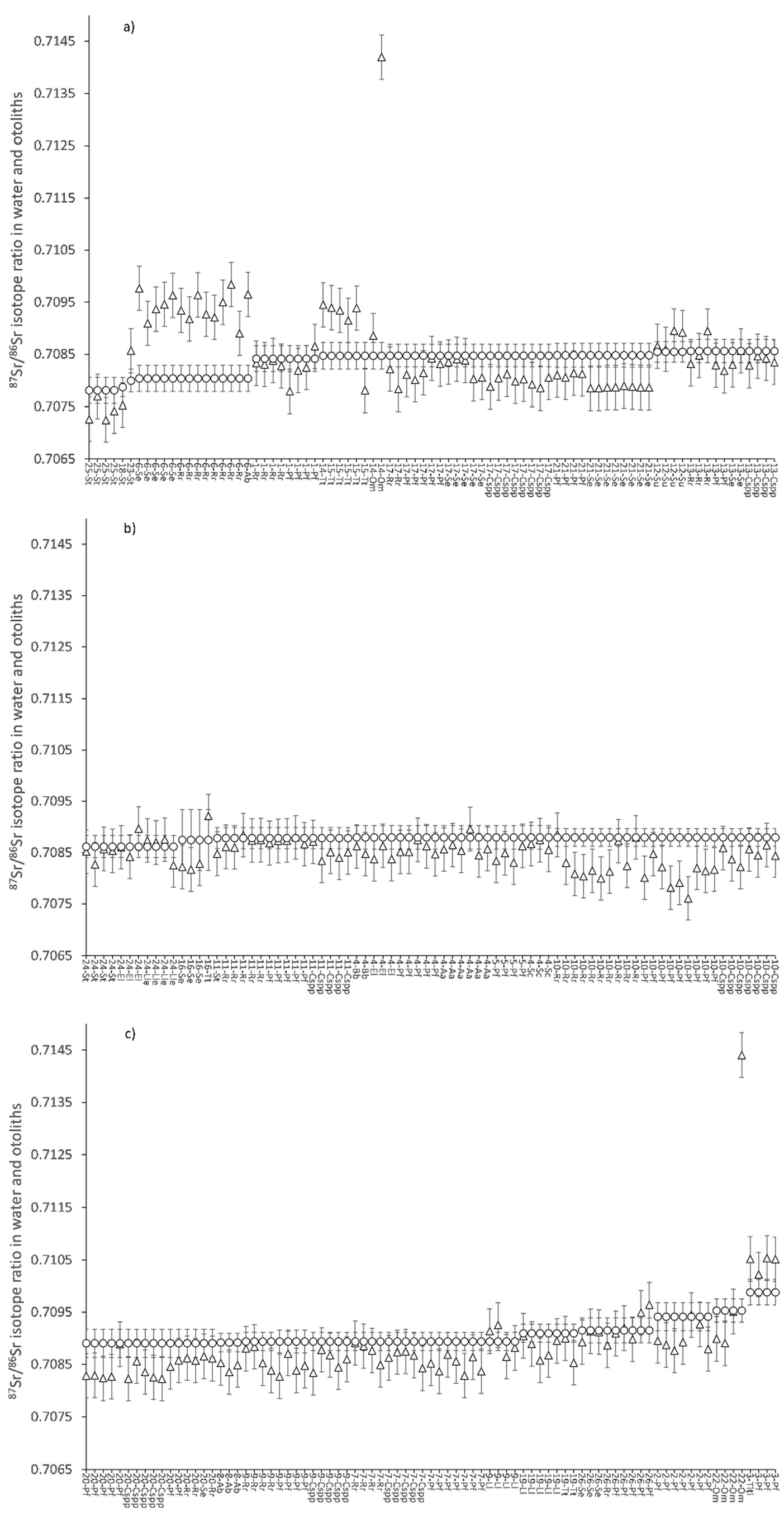
^87^Sr/^86^Sr isotope ratios of water (circles) and corresponding otolith samples (triangles) plotted ranked according to increasing ^87^Sr/^86^Sr isotope ratio of water samples with expanded uncertainties (*U*, *k*=2) for identifying transfer of fish, migration and stocking; numbers correspond to habitats described in Tab. 1 combined with fish species abbreviations (Aa=*Anguilla anguilla*, Ab=*Abramis brama*, Bb=*Barbus barbus*, Cspp=*Coregonus spp,* El=*Esox lucius*, Pf=*Perca fluviatilis*, Lle=*Leucisus leuciscus*, Ll= *Lota lota*, Om=*Ocorhynchus mykiss*, Rr=*Rutilus rutilus*, , Sc=*Squalius cephalus*, Se=*Scardinius erythrophthalmus*, St=*Salmo trutta*, Su=*Salvelinus umbla*, Tt=*Thymallus thymallus,* Titi=*Tinca tinca*).

The ^87^Sr/^86^Sr isotope ratios of 215 out of 246 otoliths overlapped within the limits of uncertainty with ^87^Sr/^86^Sr isotope ratios of water sampled from the habitats of origin. This suggested that they could be considered as having been locally inhabiting these habitats (Fig. 3). However, the ^87^Sr/^86^Sr isotope values in otoliths of 31 fish (12.6 %) did not match the ^87^Sr/^86^Sr isotope values of the water of origin within the limits of uncertainty (Fig. 3), thus these fish were considered as transferred, stocked or migrating from elsewhere to the capture place.

In particular, it was found that most fish from sampling site 6 (Baggerweiher Übersee) (eight roach *R. rutilus*, five rudd *S. erythrophthalmus* and one common bream *A. brama)* showed significantly higher ^87^Sr/^86^Sr isotope ratios as compared to the value of the water (Fig. 3a). After further investigation by contacting the responsible fishery managers, it could be proven, that this lake was stocked with cyprinids from other sources (e.g. another pond, not existing at the time of the data analysis any more) as prey for predatory fish such as *Sander lucioperca*. Hence, these fish were being considered as transferred to this lake and removed from further data analysis. Next at River Inn at habitat 14 (Inn at Nussdorf) and habitat 15 (Inn at Neubeuern) five individuals of *T. thymallus* showed significantly deviating ^87^Sr/^86^Sr isotope ratios as compared to the values of the water (Fig. 3a). As *T. thymallus* is known as a species migrating upstream for spawning, the otolith values were compared to ^87^Sr/^86^Sr isotope ratios from a downstream site near the mouth of the River Inn (about 100 km downstream) where samples were collected during an earlier study (Kendlbacher, 2013). As the ^87^Sr/^86^Sr isotope ratio of 0.70948 (± 0.00026 *U*, *k*=2) from this site matched the otolith data very well, it was concluded that these fish might have migrated upstream from a lower section of the river. Therefore, they assigned to this habitat for further analysis. One rainbow trout (*Oncorhynchus mykiss*) was obviously stocked to habitat 14 (Inn at Nussdorf, Fig. 3a), and another one from habitat 22 (Tiroler Achen, Fig. 3b) also showed a significantly higher ^87^Sr/^86^Sr isotope ratio as compared to the value of the water (similar to the other rainbow trout`s values; Fig. 3a), why it was concluded it was also stocked. Both individuals were removed from further analyses. Significantly lower ^87^Sr/^86^Sr isotope values as compared to the values of the water samples were also found in five *R. rutilus* and six *P. fluviatilis* otoliths from Lake Chiemsee near Prien (habitat 10; Fig. 3b). These values better matched with the ^87^Sr/^86^Sr isotope values of the River Prien (habitat 18, associated to the SIG with the lowest ^87^Sr/^86^Sr isotope ratio values) entering Lake Chiemsee, thus it was concluded that these fish were also using the bay area at the mouth of the River Prien as habitat. For further analysis of these fish were assigned to habitat 18 (river Prien). Thus, for the analysis of the relation between water Sr/Ca and the otolith Sr elemental mass fraction 230 otoliths were finally available.

### 3.3 Relation between Sr/Ca ratio in water and the Sr elemental mass fraction in otoliths

A significant positive correlation between Sr/Ca in water and the Sr mass fraction in otoliths was found for *Coregonus spp* (*n*=39, *r*_s_ = 760, *P* = 0.000), *E. lucius* (*n*=6, *r*_s_ = 878, *P* = 0.021), *Lota lota* (*n*=9, *r*_s_ = 866, *P* = 0.003), *Perca fluviatilis* (*n*=71, *r*_s_ = 835, *P* = 0.000), *R. rutilus* (*n*=36, *r*_s_ = 812, *P* = 0.000), *S. erythrophthalmus* (*n*=22, *r*_s_ = 710, *P* = 0.000) and *T. thymallus* (*n*=9, *r*_s_ = 743, *P* = 0.022). No correlation could be found for *S. trutta* (*n*=11, *r*_s_ = 334, *P* = 0.316) and *O. mykiss* (*n*=4, *r*_s_ = −258, *P* = 0.742). For the other species, not enough samples from different habitats to conduct the analysis were available. When analyzing the relationship between the Sr/Ca in water and the Sr mass fraction in otoliths of all cyprinids together, the correlation remained significant (*n*=71, *r*_s_ = 783, *P* = 0.000), while combining all salmonids (except *T. thymallus* and those to be considered as stocked) still yielded no significant correlation (*n*=19, *r*_s_ = 111, *P* = 0.650).

Regression analysis of Sr/Ca ratios in water and Sr mass fractions in otoliths highlighted the established correlations (Tab. 4, Fig. A.3). It became evident, that the relation in *S. erythrophthalmus* was only expressed weakly (Fig. A.3) although the correlation was highly significant. This also influenced the relation when all cyprinids were analyzed together (Fig. A.3). The data plot combined with the regression analysis also highlighted the lack of a correlation for all salmonids combined (Fig. A.3).

**Table 4:** Regression formulas describing the relationship between the Sr/Ca ratio in water and the Sr mass fraction (μg g^−1^) in otoliths of the respective species and family, with discrimination factor *D* ± standard deviation (SD) between water Sr/Ca and otolith Sr/Ca; for Salmonids *T. thymallus* – Tt was excluded; significant regressions with an adjusted *R*^2^ > 0.7 are marked in bold.

| Species/family | *n* | Regression formula | Adjusted $R^2$ | *P* | *D* | ± SD |
| --- | --- | --- | --- | --- | --- | --- |
| *Abramis brama* | 3 | - | - | - | 0.497 | 0.016 |
| *Anguilla anguilla* | 6 | - | - | - | 0.251 | 0.094 |
| *Barbus barbus* | 2 | - | - | - | 0.352 | 0.050 |
| ***Coregonus spp.*** | **39** | **$y = 175913x - 30.384$** | **0.899** | **0.000** | **0.420** | **0.057** |
| **Cyprinids** | **71** | **$y = 136057x + 172.129$** | **0.721** | **0.000** | **0.465** | **0.136** |
| *Esox lucius* | 6 | - | - | - | 0.265 | 0.050 |
| *Leuciscus leuciscus* | 4 | - | - | - | 0.480 | 0.080 |
| ***Lota lota*** | **9** | **$y = 168121x - 213.818$** | **0.761** | **0.001** | **0.327** | **0.032** |
| *Oncorhynchus mykiss* | 4 | - | - | - | 0.217 | 0.060 |
| ***Perca fluviatilis*** | **71** | **$y = 94822x + 59.134$** | **0.720** | **0.000** | **0.275** | **0.069** |
| ***Rutilus rutilus*** | **36** | **$y = 143814x + 167.778$** | **0.825** | **0.000** | **0.496** | **0.145** |
| *Scardinius erythrophthalmus* | 22 | $y = 15781x + 450.785$ | 0.081 | 0.107 | 0.464 | 0.153 |
| *Salmo trutta* | 11 | $y = 186739x - 93.695$ | 0.161 | 0.122 | 0.425 | 0.201 |
| *Salvelinus umbla* | 4 | - | - | - | 0.690 | 0.071 |
| Salmonids (except Tt) | 19 | $y = 123736x + 227.713$ | 0.181 | 0.039 | 0.437 | 0.221 |
| *Squalius cephalus* | 3 | - | - | - | 0.426 | 0.052 |
| ***Thymallus thymallus*** | **9** | **$y = 221504x - 526.484$** | **0.775** | **0.001** | **0.345** | **0.067** |
| *Tinca tinca* | 1 | - | - | - | 0.501 | - |

Significant differences in the Sr mass fractions in otoliths of different species from the same habitats were found between *P. fluviatilis* and especially cyprinids such as *R. rutilus* and *Coregonus spp.* (Fig. A.4). *P. fluviatilis* otoliths of the same habitats contained on average 60 % of the Sr elemental mass fractions of *R. rutilus* and *Coregonus spp.* otoliths as determined by the regression formulas, with *x* being the Sr/Ca ratio of the water and *y* being the Sr mass fraction (μg g^−1^) in otoliths (Tab. 4). The mean discrimination factor describing the relationship between Sr/Ca ratio in water and otoliths was 0.375 (± 0.138 SD) for all 230 otoliths. Discrimination factors ranged from 0.217 (± 0.060 SD) for *O. mykiss* to 0.690 (± 0.071 SD) for *S. umbla.*

### 3.5 Discriminating fish according to their habitat cluster

After the transfer of the cluster numbers to fish otoliths, discriminant analysis yielded 100 % correctly classified individuals for *Coregonus spp.* and *T. thymallus* and 92 % for *P. fluviatilis* and Cyprinids according to their habitat cluster of origin respectively (Fig. 4–1). Salmonids (without *T. thymallus*) could be discriminated correctly by 74 % into 5 clusters (Fig. 4–2). For other species from separate family groups such as *A. anguilla*, *E. lucius*, *L. lota* not enough individuals were available from different habitats to conduct the analysis (Fig. 4–2).

**Figure 4:**
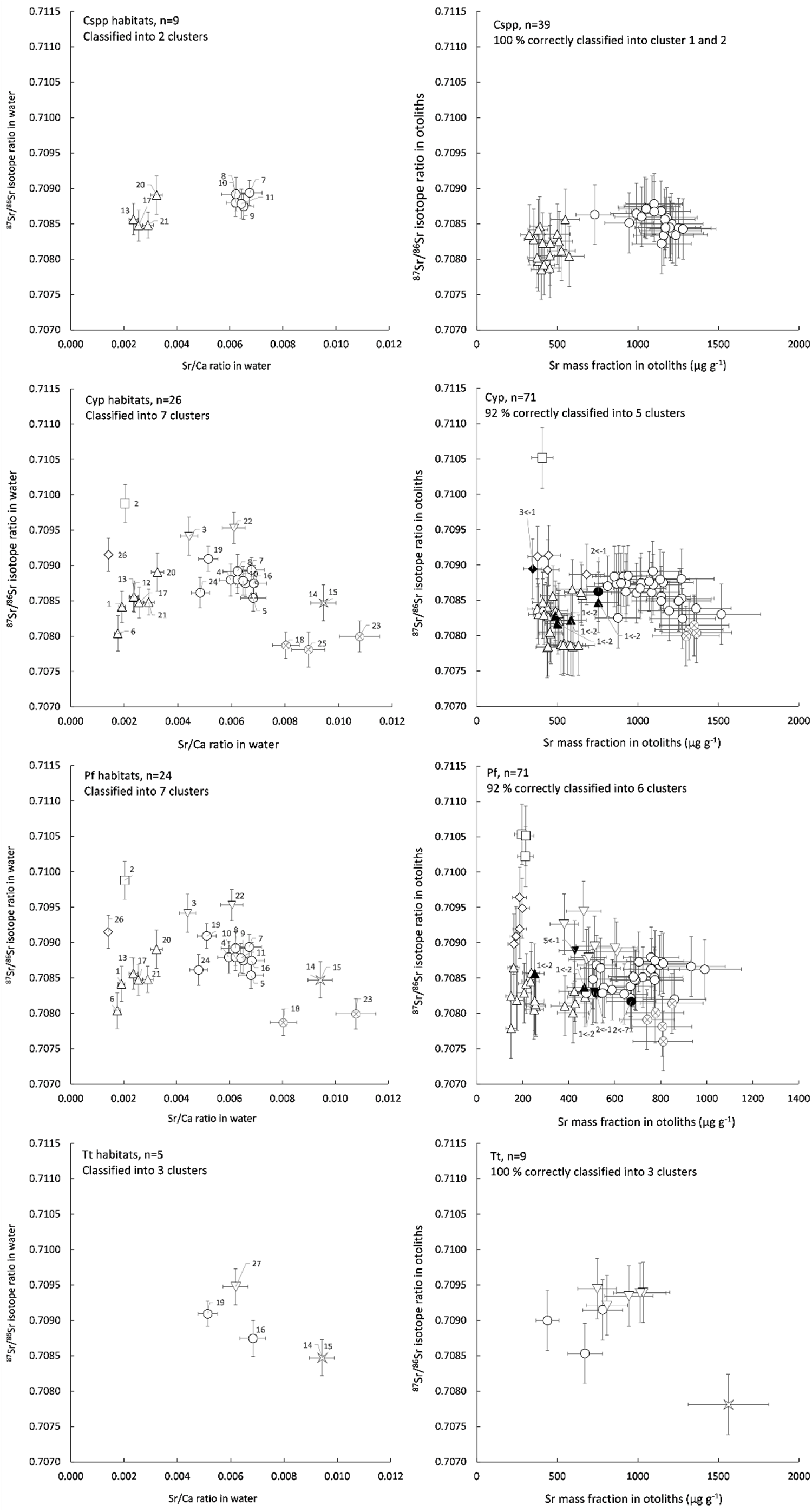
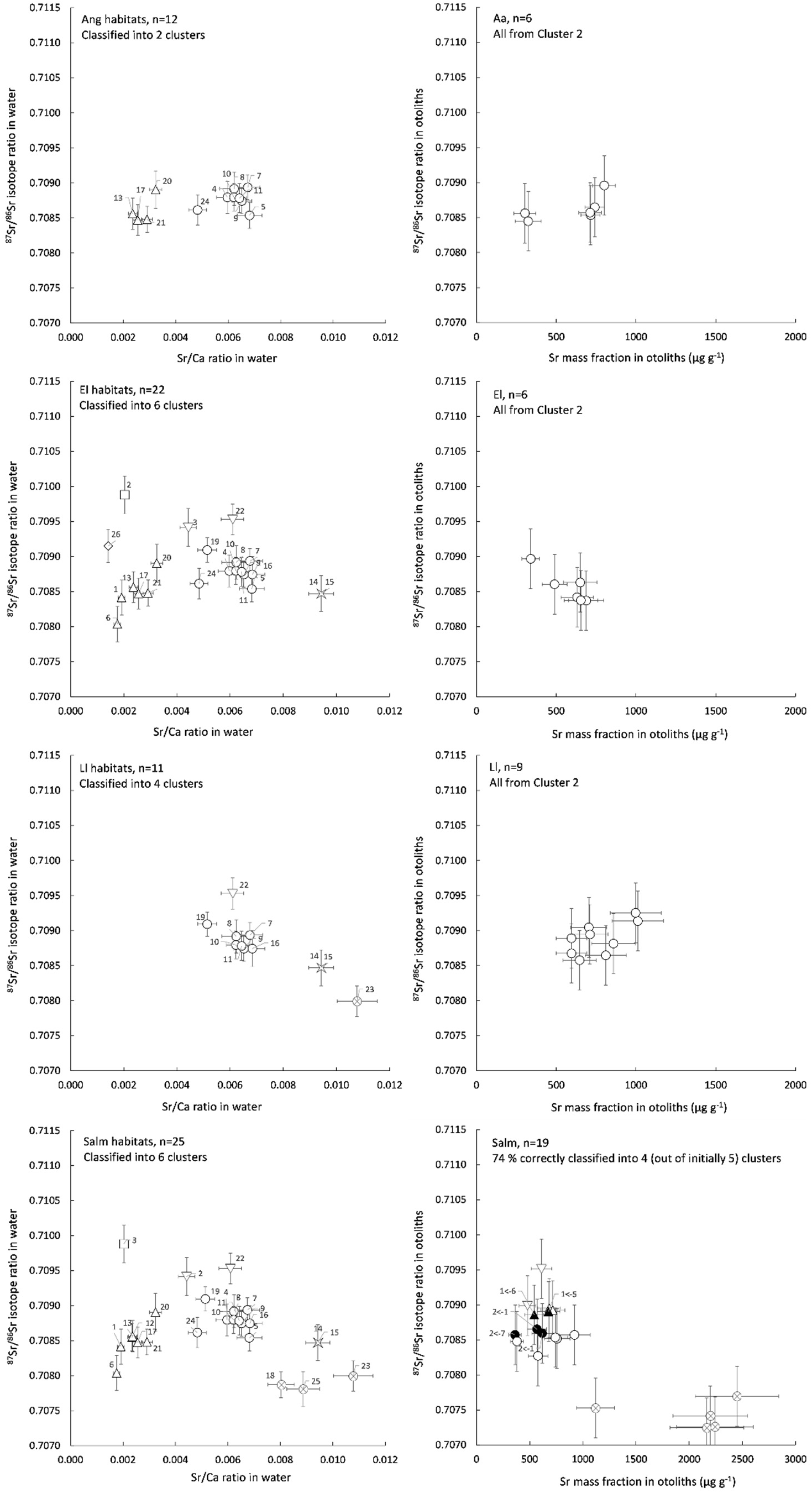
Clusters of habitats (indicated by the signatures based on Fig. 2: triangle = cluster 1, circle = cluster 2, diamond = cluster 3, square = cluster 4, flipped triangle = cluster 5, star = cluster 6, circle with cross = cluster 7) containing only those habitats where the species or family under consideration (Aa=*Anguilla anguilla*, Cspp=*Coregonus spp,* Cyp=cyprinid*s* El=*Esox lucius*, Pf=*Perca fluviatilis*, Ll= *Lota lota*, Salm= salmonids, Tt=*Thymallus thymallus*) was expected (left) and results of the discrimination analysis (right), displaying correctly (empty signatures) and incorrectly (filled signatures, with information on original cluster) classified samples; in addition the total number of individuals, the number of clusters where fish have been caught from, and the information on the percentage of correctly classified individuals are given; error bars represent expanded uncertainties (*U*, *k*=2)

## 4. Discussion

In this study, the ^87^Sr/^86^Sr isotope ratio in combination with the Sr mass fraction in fish otoliths was comprehensively evaluated as a potential complementary fishery management tool in an Alpine foreland with limited geological variability. For the application of Sr tags in otoliths to differentiate fish according to their origin, a sufficient variability of the ^87^Sr/^86^Sr isotope ratios within the study area is required, which is mainly driven by the underlying geology. A significant variation in water chemistry at spatial scales ranging from less than 1 to several hundreds of kilometers is possible (Elsdon et al., 2008). In the present study, the geology around lake Chiemsee (radius of about 50 km) displayed limited variability. Nonetheless, three clearly distinguishable strontium isotope groups *sensu* Brennan et al. (2015b) could be differentiated by the ^87^Sr/^86^Sr isotope ratios and their expanded uncertainties (*U*, *k*=2). Most of the ^87^Sr/^86^Sr isotope ratios ranged between 0.7085 and 0.7900 and were represented as one SIG containing 13 habitats, while the lowest measured ^87^Sr/^86^Sr value was 0.7078 and the highest ^87^Sr/^86^Sr value 0.7099. The Sr/Ca ratio proved to be particularly useful to further differentiate the SIGs into seven habitat clusters, as water from different habitats showed significantly different Sr/Ca ratios. Building habitat clusters with similar chemical water characteristics is a recommended approach for a conservative classification of fish according to their origin given measurement uncertainties and natural variation (Wells et al., 2003; Zitek et al., 2010). As clustering as a standard tool for biologists is known to be an imperfect process (Kern et al., 2017), assignment of weights to variables according to subject matter knowledge for clustering (Kaufman and Rousseeuw, 2005) or the combination with visual inspection and refinement using contextual data (Kern et al., 2017) have been suggested. Within this study, the relevance of clearly distinguishable SIGs together with the variation of the Sr/Ca ratio and the expanded combined standard uncertainties were used as contextual data for improving the grouping of water data into clusters. Finally, all Lake Chiemsee habitats (cluster 2) could be clearly differentiated from all other smaller lakes around (cluster 1) and single lacustrine habitats (clusters 3, 4 and 5) (Fig. 4). Clusters 6 and 7 only contained riverine habitats, with cluster 6 representing two sites of the River Inn, one of the two large rivers crossing the study area in Germany. The other larger river, the River Salzach was clustered together with a more downstream section of the River Inn together with Lake Chiemsee (cluster No. 2). Both rivers have their sources in Austria and cut through igneous, metamorphic and sedimentary rock in different degrees along their course (Janoschek and Matura, 1980). The ^87^Sr/^86^Sr isotope ratio of the River Inn increases during its downstream course, with the River Salzach being even higher in ^87^Sr/^86^Sr isotope ratio at the study site. Later, the River Salzach joins the River Inn and contributes to its elevated ^87^Sr/^86^Sr isotope ratio at its mouth (Kendlbacher, 2013). Cluster 7 contains the rivers Prien, Weißach and Traun, with the River Prien entering Lake Chiemsee, characterized by the lowest ^87^Sr/^86^Sr isotope ratios and combined in one SIG. Prien and Traun, but also the much smaller Weißach, have their sources in sedimentary limestone dominated geological formations (Fig. 1), explaining the low ^87^Sr/^86^Sr isotope ratio (Faure and Mensing, 2005). The results prove that hypothesis (1) “differences in the ^87^Sr/^86^Sr isotope and Sr/Ca ratios between the investigated water bodies exist and can be used to determine useful habitat clusters” can therefore be supported. However, there will always be some uncertainty involved in this approach, especially when considering expanded measurement uncertainties. Thus, the exploration of expected differences in water chemistry with specific questions in mind is recommended before conducting otolith studies (Wells et al., 2003; Humston and Harbor, 2006).

In total 31 out of 246 analyzed otoliths were associated to fish that have been transferred, stocked or which might have migrated using the comparison between ^87^Sr/^86^Sr isotope ratios in water and otoliths under consideration of the expanded uncertainties (*U*, *k*=2). In total two *O. mykiss* were considered as stocked and 13 cyprinids (eight roach *R. rutilus*, five rudd *S. erythrophthalmus)* as transferred to a lake as feed for *S. lucioperca* while five grayling *T. thymallus* were considered as having moved upstream to the capture site and five roach *R. rutilus* and six European perch *P. fluviatilis* from lake Chiemsee were considered using the bay at the mouth of the River Prien into Lake Chiemsee as a habitat. Salmonids, such as *O. mykiss*, are the most stocked fish species for fishery purpose in Europe. Thus, it can be expected that these fish stem for fish farms when significant differences in the ^87^Sr/^86^Sr isotope ratios between water and otoliths exist. Unfortunately, we did not have more information on potential source fish farms for these fish, why they could not be matched to any specific source habitat. In general, when information on the source hatcheries is available, the ^87^Sr/^86^Sr isotope can be used as a natural tag to track back fish to their source hatchery (Zitek et al., 2010). On the other hand, European grayling *T. thymallus* are hardly stocked as adults. Therefore, deviations between the ^87^Sr/^86^Sr isotope ratio of otoliths and water were interpreted as migrations. In relation to potential source habitats for these fish, data from a site approximately 100 km downstream at the mouth of the River Inn were available from an earlier study (Kendlbacher, 2013). They could be successfully matched with the ^87^Sr/^86^Sr isotope data of the grayling otoliths supporting the hypothesis that fish stem from a downstream region. However, it could not be finally proven, from which distance the fish exactly stemmed. European grayling is supposed to perform maximum upstream migrations of about 30 km (Waidbacher and Haidvogl, 1998), but water data from this region were not available. In addition, the connectivity situation would have needed consideration, as the River Inn has several hydropower plants, with only some of them equipped with fish passes during the time of the study. This fact points towards a potential application of otolith microchemistry to indirectly assess the connectivity of a river catchment system, the effect of dams and effectivity of connectivity measures in fragmented river systems (Clarke et al., 2007; Fukushima et al., 2014). Concluding, for freshwater fish species conducting migrations over tens and potentially hundreds of km it is also necessary to look for potential source habitats in downstream sections of a river and include these sites for sampling and analysis. Depending on the question, a full coverage of the main river stem including its tributaries to construct an isotopic landscape (so called “isoscape”) can considered as one important basis for an informed application of otolith microchemistry with regard to migration and dispersal of fish in a river catchment (Muhlfeld et al., 2012).

The relationship between habitat Sr/Ca ratio and the Sr mass fractions in otoliths was only analysed for fish where a clear direct relationship between the habitat and the fish could be assumed. Typically, as fish take up Sr from water by its relative availability in relation to Ca (Campana, 1999), several studies related the Sr/Ca ratio in water successfully to the associated changes in otolith Sr mass fractions by regressions (Wells et al., 2003; Zeigler and Whitledge, 2010; Strohm et al., 2017). The current study supports these findings by showing also a clear positive trend between the Sr/Ca ratios in water and the Sr mass fractions in otoliths. The regressions were determined for different species and families, and clear species-specific trends could be identified. *R. rutilus* and *Coregonus spp.* showed the highest uptake of Sr, while in comparison other species such as *P. fluviatilis* from the same water body took up only about 60 % of Sr. Species specific differences in the uptake of Sr from the environment have been documented for marine (Swearer et al., 2003) and freshwater species (Friedrich and Halden, 2008). Interspecific variation in otolith microchemistry in fish from same habitats is generally expected to be more expressed in phylogenetically or ecologically more distantly related species (Swearer et al., 2003). However, for typical European freshwater species information on interspecific differences are rare. For example Lenaz et al. (2006) reported interspecific differences of Sr concentrations in lapilli of three different typical European freshwater fish species like the lacustrine species common carp *Cyprinus carpio*, tench *T. tinca* and a rheophilous cyprinid, nase *Chondrostoma nasus*, with the Sr mass fraction in otoliths of the latter being significantly lower as compared to the two others.

As compared to cyprinids, *P. fluviatilis* and *Coregonus spp*, salmonids only showed a weak relationship between Sr/Ca ratios in water and the Sr elemental mass fraction in otoliths in this study. This can be explained by the fact, that salmonids are regularly stocked as adults for fishery purposes. Although it was tried to identify stocked fish by deviating ^87^Sr/^86^Sr isotope ratios, still the opportunity remains, that fish with similar ^87^Sr/^86^Sr isotope ratios in otoliths could also have been stocked from a hatchery displaying the same ^87^Sr/^86^Sr isotope ratio but having a different Sr/Ca ratio. Stocking hereby significantly can impair the identification of the source of the fish. However, when including the whole life history reflected in an otolith by varying ^87^Sr/^86^Sr isotope ratios and Sr elemental mass fractions by cross-sectional line scans as shown by Zitek et al. (2010), this problem could be targeted.

To further characterize the species specific differences in Sr uptake from water in the same environment, partition coefficients as a measure for species specific Sr discrimination were calculated as discrimination D_Sr:Ca_ = (Sr:Ca)_otolith_/(Sr:Ca)_water_ (Campana, 1999). For this purpose elemental mass fractions in otoliths had to be first converted to Sr/Ca mass fraction ratios (μg g^−1^), which was done using the stoichiometric concentration of Ca in aragonite (Whitledge et al., 2019).

Although partition factors between water and otolith Sr for selected non-European freshwater salmonids exist (Wells et al., 2003; Muhlfeld et al., 2012; Stewart et al., 2021) only individual data sets for comparable typical European freshwater fish species are currently available (Melancon et al., 2009). Partition factors documented in this study were 0.420 ± 0.057 SD for *Coregonus spp.*, 0.465 ± 0.136 SD for cyprinids, 0.275 ± 0.069 SD for *P. fluviatilis,* and 0.496 ± 0.145 SD for *R. rutilus*. In comparison Melancon et al. (2009) documented a discrimination factor of 0.35 for burbot *L. lota,* which is comparable to the value of 0.327 ± 0.032 SD found for *Lota lota* in the present study. Clarke et al. (2007) described a partition factor of 0.37 for arctic grayling *Thymallus arcticus* which is comparable to the value of 0.345 ± 0.067 SD documented for *T. thymallus* in this study. The discrimination factors for typical non-European salmonids documented in literature range from 0.28 for Lake trout *Salvelinus namaicush* (Melancon et al., 2009), 0.285 ± 0.047 SD for westslope cutthroat trout (*Oncorhynchus clarkii lewisi*) (Muhlfeld et al., 2012), 0.29 for nonindigenous lake trout in the Yellowstone park (*Salvelinus namaycush*) (Stewart et al., 2021) to 0.4 for field-sampled westslope cutthroat trout *O. clarki lewisi* (Wells et al., 2003). These lower values of around 0.28 almost match the value of *O. mykiss* otoliths (*n*=4) of 0.217 ± 0.060 SD in this study. *O. mykiss* is a non-native species in Europe and was introduced from USA in the late 19^th^ century. Nowadays this species has established self-sustaining populations in many European regions (Stanković et al., 2015). Considering the above-described facts, hypothesis (2), that “species differ regarding the uptake of Sr from the environment”, can be accepted. However, still other factors, that might affect the uptake ratio of Sr in relation to the environmental availability, need consideration when conducting otolith microchemistry studies. Besides the documented species-specific effects, intraspecific differences in the uptake ratio of Sr caused by environmental conditions in habitats or simply by the age/size affecting the protein synthesis in relation to the crystallization rate of the otolith might be also existing (Campana, 1999). For example, an increasing trend of the Sr/Ca ratio with fish size independent of environmental variability indicating difference in elemental uptake with age was documented for a salmonid species only recently (Stewart et al., 2021). Basically, discrimination of elements can occur at three physiological barriers in freshwater fish, from water to gill, from blood to endolymph liquid and finally from endolymph to the otolith via the crystallization process (Campana, 1999; Melancon et al., 2009).

Clustering habitats with similar chemical characteristics is an important first step to identify the possibilities to discriminate fish according to their source and origin (Wells et al., 2003; Zitek et al., 2010). In the present study, seven water clusters with distinct^87^Sr/^86^Sr isotope and Sr/Ca ratios at a meaningful scale for management purposes could be formed. In a next step, based on the information of potential occurrence of fish in specific habitats, the habitat clusters were further reduced on a taxon-specific level. This yielded for example two distinct clusters, where *Coregonus spp*. might be expected, while for cyprinids or *P. fluviatilis* all seven clusters were of potential relevance. After the transfer of the cluster numbers to fish otoliths, individuals of *Coregonus spp.* and *T. thymallus* could be correctly classified by 100 %, while for perch *P. fluviatilis* and for cyprinids the correct classification rate was 92 %. Salmonids (without *T. thymallus*) were classified correctly by 74 %. The lack of individuals from different habitats for other species such as *A. anguilla*, *E. lucius*, *L. lota* did not allow for this analysis. Because of the different levels of aggregation of habitats, it is hard to compare the classification success rates of this study to international studies. However, it has been recognized that even when no site specific differentiation of fish can be achieved, classification into habitat types and clusters at a higher level of integration still can be informative for fisheries management (Radigan et al., 2018c). Furthermore Radigan et al. (2018c) noted, that 80 % correct classifications can be seen as threshold for reliable otolith chemistry studies. This value can be used to support the stepwise aggregation of habitats to a specific scale of analysis to achieve potential meaningful results for fishery management. Hypothesis (3), that “otoliths could be assigned to their habitat clusters of origin with a high probability when potential species-specific differences are considered” can be therefore accepted for *Coregonus spp.*, Cyprinids, *T. thymallus* and *P. fluviatilis* when 80 % correct assignments to a cluster are defined as a minimum value for reliable otolith microchemistry studies. As mentioned earlier, if otolith microchemistry is considered as a potential tool for fishery management, it is relevant to start with an understanding of how distinct water chemistry among habitats at the scales of interest is, and potentially consider the integration of otolith chemistry with complementary techniques (Radigan et al., 2018c). Especially the focus on specific questions such as stocking, source of fish, migration and dispersal etc. will influence the definition of the scale of the study and sampling design, the potential usefulness of aggregating habitats into clusters and selection of complementary methods. Complementary techniques that have been suggested for larger scales are e.g. genetics or mark-recapture experiments, while for smaller scales e.g. radio-telemetry or other isotopic systems in muscles could be used (Pracheil et al., 2014). Furthermore, there are indications, that also interpopulation in otolith shape could support the discrimination of otoliths according to their origin (Souza et al., 2020; Zitek, unpublished data, this study). Summarizing, species and habitat characteristics and question under study are the major considerations for otolith chemistry studies that are useful in terms of fisheries management (Radigan et al., 2018a).

## 5. Conclusion

The results of the present study show that otolith microchemistry can be applied as a potential fisheries management tool for various typical European freshwater fish species even at a relatively small a spatial scale in an Alpine foreland with limited geological variability. It provides a substantial basis for hypothesis building and future applications of the of ^87^Sr/^86^Sr isotope ratio and Sr/Ca ratio as a fishery management tool for. Researchers trying to determine the origin of fish based on otolith data should be aware of the fact that stocking of fish for fishery purposes, potential transfer and migration of fish frequently occur and should be considered in the design of the study. The expected differences in water chemistry will affect the potential conclusions that could be drawn from the data for a specific question and species. Therefore, a preliminary assessment of the potential differences in water chemistry at the scale of the study are recommended, as well as the collection of data on the distribution and natural migration pathways of different fish species under study. Potential future applications of otolith microchemistry for the management of European freshwater species are e.g. the assessment of seasonal habitat use, migrations and dispersal including the indirect assessment of the passability of artificial barriers for fish, the identification and discrimination of the most relevant spawning and rearing habitats for self-sustaining populations and the contribution of stocked fish to natural populations. Moreover, the prey of piscivores can be retrospectively provenanced to identify frequented foraging habitats. In the area of food fraud, regionally produced fishery and aquaculture products can be authenticated and mislabeling of these products can be identified. Future studies should focus on a better understanding of the relation between environmental concentrations of elements like Sr, environmental factors and fish growth/size and otolith microchemistry for different typical European freshwater species. Complementary methods like the analysis of the otolith shape that could contribute to an improved discrimination of fish according to their habitat of origin could be considered.

## Acknowledgements

This study was funded by the Austrian Science Fund (FWF), project number P24059, entitled “The feeding ecology of the Great Cormorant.” We also acknowledge financial support through grants from Hypo Tirol Bank, Swarovski, and PhD scholarships (University of Innsbruck) for JO and BT, and a research scholarship for Austrian graduates of the University Innsbruck awarded to BT. We thank H. Schaber, T. Lex, I. Wallner, F. Kirchmeier, F.Kirmeier, J. Kraller, B. Graßl, J. Kern, J. Spatzenegger, J. Hartl, J. Haiker, K. Kreissig, F. Göpfert, and P. Huber for providing fish or giving us permission to catch them per electric fishing. We also thank the water guard of Lake Chiemsee for their service regarding the collection of water samples at Lake Chiemsee as well as the fish farms Eulenau, Jäckle, Kreissnig and Weiß for their permission to take water samples at their ponds. We also thank the Bavarian state agency for environment for providing digital maps of the study area. Final analysis and interpretation of otolith data was also supported by the COMET-K1 competence centre FFoQSI, funded by the Austrian ministries BMVIT, BMDW and the Austrian provinces Niederoesterreich, Upper Austria and Vienna within the scope of COMET - Competence Centers for Excellent Technologies. The programme COMET is handled by the Austrian Research Promotion Agency FFG. The strategic objectives of COMET are: developing new expertise by initiating and supporting long-term research co-operations between science and industry in toplevel research, and establishing and securing the technological leadership of companies. By advancing and bundling existing strengths and by integrating international research expertise Austria is to be strengthened as a research location for the long term.

## Author contributions

**Andreas Zitek:** Study Conceptualization, Otolith Preparation for Laser Ablation ICP-MS, Data Analysis and Statistics, Visualisation, Writing-Original Draft, Writing-Reviewing and Editing; **Johannes Oehm:** Otolith and Water Sample Collection, Otolith Extraction and Preparation, Geodata Collection and Provision, Writing-Reviewing and Editing; **Michael Schober:** Laser Ablation ICP-MS Measurements, Data Evaluation; **Anastassiya Tchaikovsky:** Water Sample Preparation, ICP-MS Measurements, Data Evaluation, Writing-Reviewing and Editing; **Johanna Irrgeher:** ICP-MS Measurements, Data Evaluation, Writing-Reviewing and Editing, Supervision; **Anika Retzmann:** Laser Ablation ICP-MS, Data Evaluation, Writing-Reviewing and Editing; **Bettina Thalinger:** Otolith and Water Sample Collection, Geodata Collection and Provision, Writing-Reviewing and Editing; **Michael Traugott:** Study Conceptualization, Writing-Reviewing and Editing, Supervision; **Thomas Prohaska:** Study Conceptualization, ICP-MS Method Development, Data Evaluation, Writing-Reviewing and Editing, Supervision.

## Notes

### Competing Interest Statement

The authors have declared no competing interest.

