## Supplementary table and figures for "Evaluating ^87^Sr/^86^Sr isotope ratios and Sr mass fractions in otoliths of different European freshwater fish species as fishery management tool in an Alpine foreland with limited geological variability"

Table A.1: Sampled water bodies, type, location with geographic coordinates and dates of water and fish samples; for six water bodies (*) the exact location of the fish samples was not available as these fish were captured by fishermen within the investigated lakes, or in one occasion, somewhere along the lower course of the River Prien.

| No | Water Body | Type | Location | Coordinates water sample | Date water sample | Coordinates fish sample | Date fish sample |
|---|---|---|---|---|---|---|---|
| 1 | Abtsdorfer See | Lake | Laufen | 47°54'52.3"N 12°54'21.5"E | 28.10.2014 | 47°54'52.3"N 12°54'21.5"E | 01.09.2013 |
| 2 | Altwasser Osterbuchberg | Oxbow | Grabenstätt | 47°48'34.6"N 12°30'10.6"E | 28.10.2014 | 47°48'30.2"N 12°30'19.4"E | 23.01.2014 |
| 3 | Almfischerweiher | Pond | Übersee | 47°48'56.8"N 12°29'57.8"E | 21.11.2012 | 47°48'53.5"N 12°29'53.3"E | 23.01.2014 |
| 4 | Alz upstream Traun entry | River | Altenmarkt | 48°00'13.4"N 12°32'00.3"E | 21.11.2012 | 47°59'53.8"N 12°31'19.9"E | 05.12.2013 |
| 5 | Alz downstream Traun entry | River | Trostberg (downstream Altenmarkt) | 48°01'50.2"N 12°33'43.9"E | 10.03.2011 | 48°01'50.2"N 12°33'43.9"E | Autum 2013 |
| 6 | Baggerweiher Übersee | Pond | Übersee | 47°50'52.5"N 12°29'12.4"E | 21.11.2012 | 47°50'53.5"N 12°29'08.7"E | 23.01.2014 |
| 7 | Chiemsee | Lake | Chieming | 47°53'09.2"N 12°30'19.5"E | 03.09.2013 | 47°52'59.3"N 12°31'14.8"E | Summer 2013 |
| 8 | Chiemsee | Lake | Felden | 47°50'28.3"N 12°23'12.6"E | 03.09.2013 | 47°50'29.2"N 12°23'06.2"E | 26.07.2013 |
| 9 | Chiemsee | Lake | Fraueninsel | 47°52'15.4"N 12°25'50.5"E | 03.09.2013 | 47°52'10.4"N 12°25'49.1"E | Summer 2013 |
| 10 | Chiemsee | Lake | Prien | 47°51'59.8"N 12°22'15.1"E | 03.09.2013 | 47°51'59.8"N 12°22'15.1"E | Summer 2013 |
| 11 | Chiemsee | Lake | Seebruck | 47°55'17.1"N 12°28'27.4"E | 03.09.2013 | 47°55'37.2"N 12°29'00.7"E | Summer 2013 |

| | | | | | | | |
|---|---|---|---|---|---|---|---|
| 12 | Fischzucht Kreissig | Fish Farm | Amerang | 47°58'37.1"N 12°17'31.9"E | 21.11.2012 | 47°58'37.1"N 12°17'31.9"E | Spring 2014 |
| 13 | Hartsee | Lake | Eggstätt | 47°55'45.9"N 12°22'26.1"E | 21.11.2012 | * | August 2013 |
| 14 | Inn Rosenheim upstream Mangfall entry | River | Nussdorf | 47°51'31.3"N 12°08'13.3"E | 08.02.2013 | 47°44'36.7"N 12°08'05.4"E | October 2013 |
| 15 | Inn Rosenheim upstream Mangfall entry | River | Neubeuern | 47°51'31.3"N 12°08'13.3"E | 08.02.2013 | 47°46'44.4"N 12°07'41.6"E | 19.10.2013 |
| 16 | Inn upstream Wasserburg | River | Griesstätt | 47°59'42.7"N 12°09'48.3"E | 08.02.2013 | 48°03'08.2"N 12°12'38.8"E | Summer 2013 |
| 17 | Langenbürgner See | Lake | Breitbrunn | 47°53'42.1"N 12°21'38.7"E | 21.11.2012 | * | 20.08.2013 |
| 18 | Prien | River | Prien | 47°51'13.6"N 12°20'06.2"E | 03.09.2013 | * | *March 2014* |
| 19 | Salzach | River | Laufen | 47°56'22.9"N 12°56'21.9"E | 10.03.2011 | 47°54'45.1"N 12°57'18.9"E | 14.10.2013 |
| 20 | Simssee | Lake | Riedering | 47°51'27.6"N 12°13'35.2"E | 10.03.2011 | * | Summer 2013 |
| 21 | Tachinger See | Lake | Taching | 47°59'18.7"N 12°44'42.1"E | 10.032011 | * | Autum 2013 |
| 22 | Tiroler Achen | River | Unterwössen/Marquartstein | 47°44'18.0"N 12°26'42.3"E | 21.11.2012 | 47°45'09.6"N 12°27'57.9"E | 05.12.2013 |
| 23 | Traun | River | Altenmarkt | 47°51'55.2"N 12°39'05.6"E | 10.03.2011 | 47°59'46.5"N 12°32'21.9"E | 05.12.2013 |
| 24 | Überseer Bach | River | Übersee | 47°50'15.2"N 12°28'51.8"E | 21.11.2012 | 47°50'02.5"N 12°28'35.0"E | 23.01.2014 |
| 25 | Weißach | River | Grabenstätt | 47°49'37.7"N 12°30'49.1"E | 21.11.2012 | 47°48'41.6"N 12°33'15.9"E | 23.01.2014 |
| 26 | Weitsee | Lake | Schnaitsee | 48°03'31.0"N 12°22'07.0"E | 08.02.2013 | * | Summer 2013 |

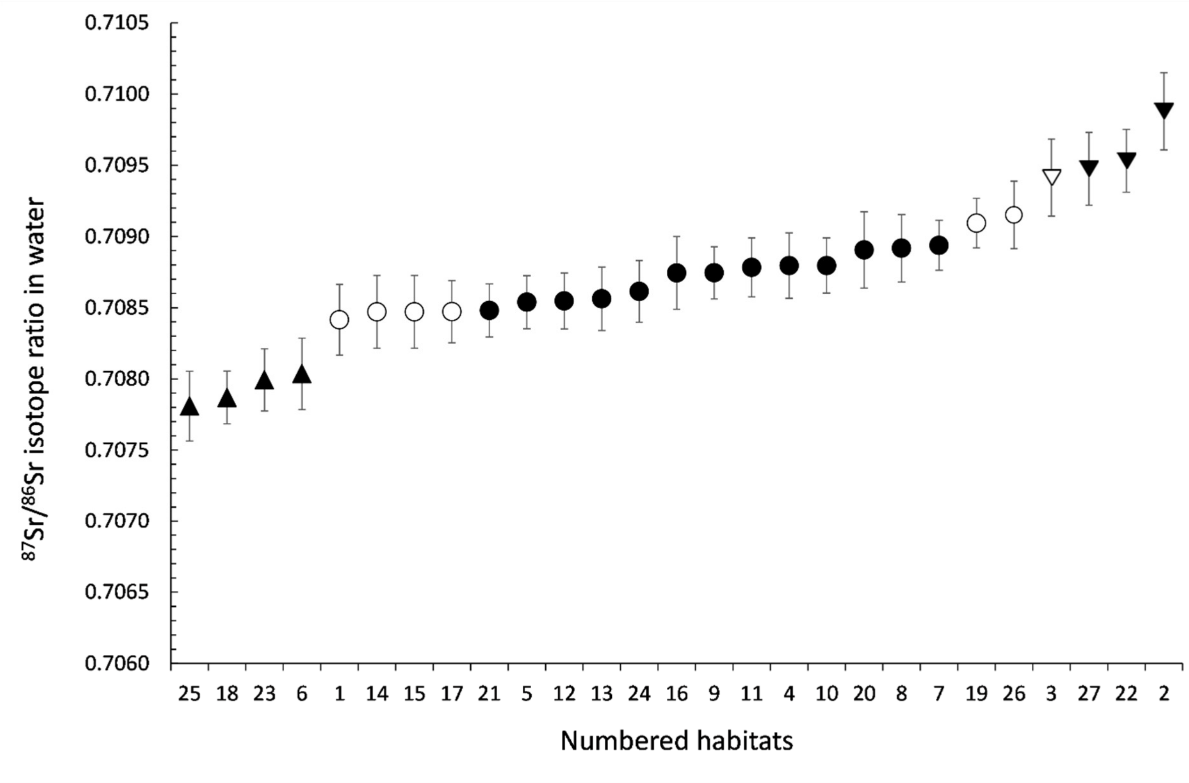


Figure A.1: Strontium isotope groups (SIGs) after Brennan et al (2015b) as determined by non-overlapping expanded uncertainties ($U$, $k$=2) in filled signatures with associated transition habitats determined by the degree of overlap of the uncertainties shown as empty signatures; habitat 27 is the mouth of the River Inn added from Kendlbacher (2013).

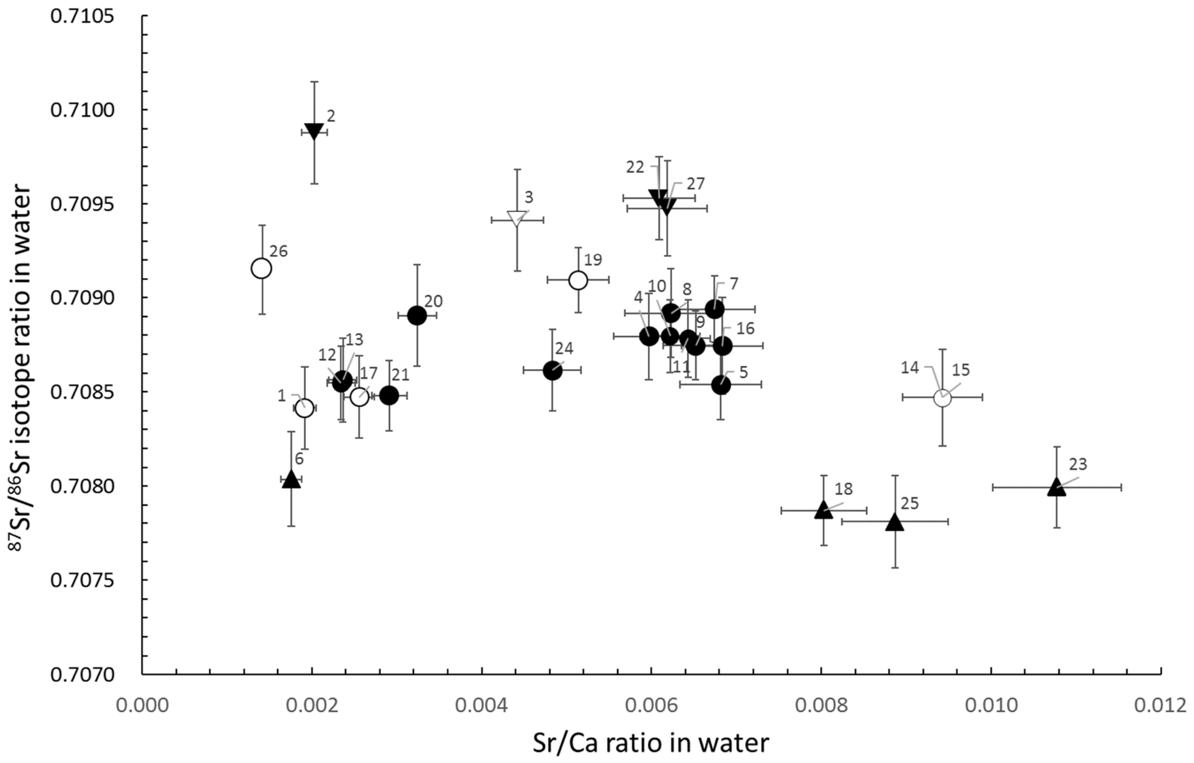


Figure A.2: Representation of the three SIGs of water samples combined with Sr/Ca values to support the discrimination of habitats into clusters; filled shapes (triangle, circle, flipped triangle) represent habitats that could be clearly be grouped into SIGs based on their not-overlapping uncertainties, open shapes represent intermediated habitats overlapping with SIG habitats in their uncertainties; habitat 27 is the mouth of the River Inn added from Kendlbacher (2013).

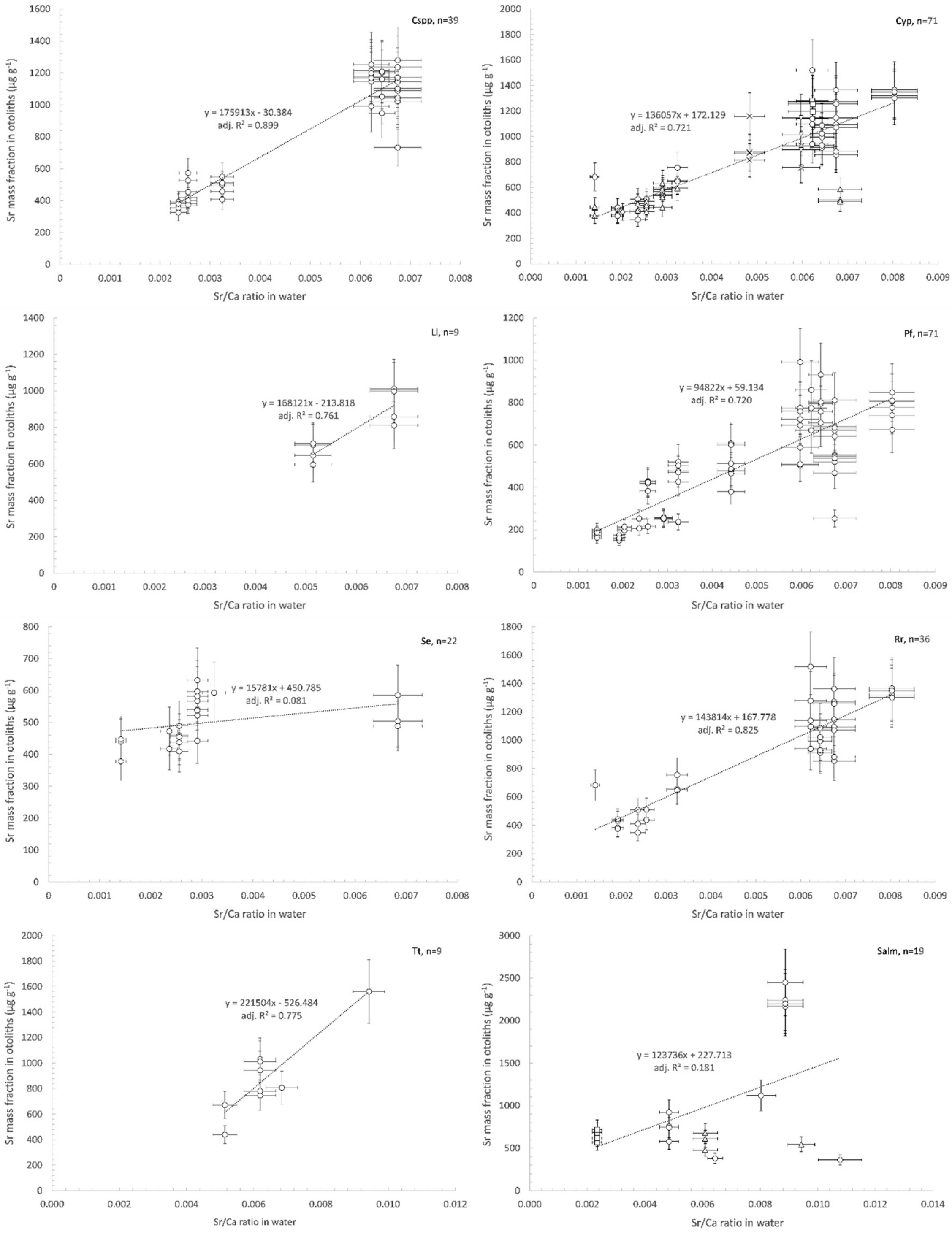


Figure A.3: Relationship between Sr/Ca ratio in water and the Sr elemental mass fraction (µg $g^{-1}$) in otoliths for roach *Rutilus rutilus* (Rr), European perch *Perca fluviatilis* (Pf), whitefish *Coregonus spp* (Cspp), rudd *Scardinius erythrophthalmus* (Se), burbot *Lota lota* (Ll), European grayling *Thymallus thymallus* (Tt), salmonids (*Salmo trutta* – circles, *Salvelinus umbla* – squares, *Oncorhynchus mykiss* - triangles) (left) and all Cyprinids (*Abramis brama* – square, *Barbus barbus* – star, *Leuciscus leuciscus* – flipped cross, *Rutilus rutilus* – circles, *Scardinius erythrophthalmus* – triangle, *Squalius cephalus* – flipped triangle, *Tinca tinca* – diamond), with expanded uncertainties (*U*, *k*=2).

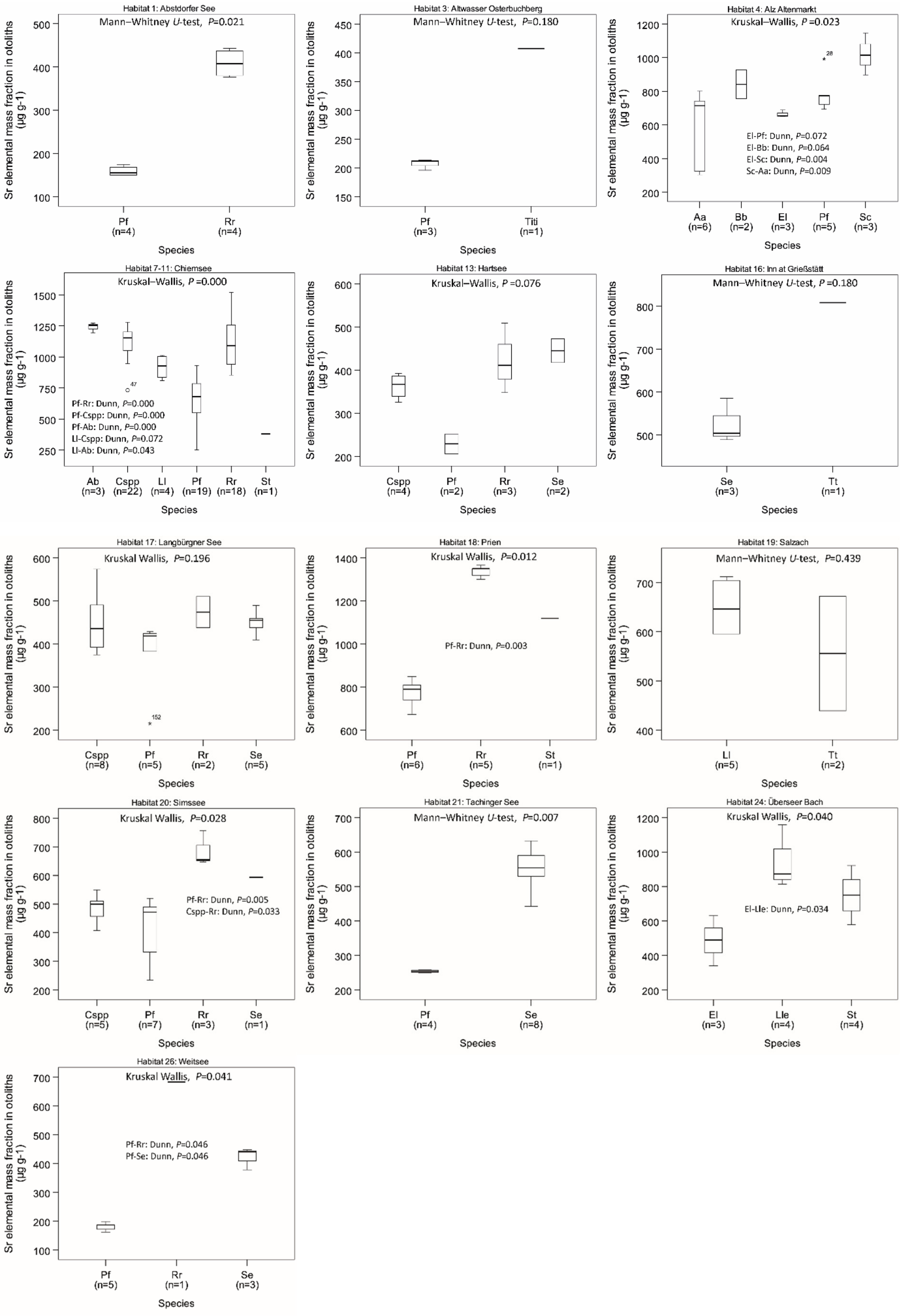

Habitat 1: Abstdorfer See
Mann–Whitney U-test, P=0.021
Pf (n=4)
Rr (n=4)
Habitat 3: Altwasser Osterbuchberg
Mann–Whitney U-test, P=0.180
Pf (n=3)
Titi (n=1)
Habitat 4: Alz Altenmarkt
Kruskal–Wallis, P =0.023
El-Pf: Dunn, P=0.072
El-Bb: Dunn, P=0.064
El-Sc: Dunn, P=0.004
Sc-Aa: Dunn, P=0.009
Aa (n=6)
Bb (n=2)
El (n=3)
Pf (n=5)
Sc (n=3)
Habitat 7-11: Chiemsee
Kruskal–Wallis, P =0.000
Pf-Rr: Dunn, P=0.000
Pf-Cspp: Dunn, P=0.000
Pf-Ab: Dunn, P=0.000
Ll-Cspp: Dunn, P=0.072
Ll-Ab: Dunn, P=0.043
Ab (n=3)
Cspp (n=22)
Ll (n=4)
Pf (n=19)
Rr (n=18)
St (n=1)
Habitat 13: Hartsee
Kruskal–Wallis, P =0.076
Cspp (n=4)
Pf (n=2)
Rr (n=3)
Se (n=2)
Habitat 16: Inn at Grießstätt
Mann–Whitney U-test, P =0.180
Se (n=3)
Tt (n=1)
Habitat 17: Langbürgner See
Kruskal Wallis, P=0.196
Cspp (n=8)
Pf (n=5)
Rr (n=2)
Se (n=5)
Habitat 18: Prien
Kruskal Wallis, P=0.012
Pf-Rr: Dunn, P=0.003
Pf (n=6)
Rr (n=5)
St (n=1)
Habitat 19: Salzach
Mann–Whitney U-test, P=0.439
Ll (n=5)
Tt (n=2)
Habitat 20: Simssee
Kruskal Wallis, P=0.028
Pf-Rr: Dunn, P=0.005
Cspp-Rr: Dunn, P=0.033
Cspp (n=5)
Pf (n=7)
Rr (n=3)
Se (n=1)
Habitat 21: Tachinger See
Mann–Whitney U-test, P=0.007
Pf (n=4)
Se (n=8)
Habitat 24: Überseer Bach
Kruskal Wallis, P=0.040
El-Lle: Dunn, P=0.034
El (n=3)
Lle (n=4)
St (n=4)
Habitat 26: Weitsee
Kruskal Wallis, P=0.041
Pf-Rr: Dunn, P=0.046
Pf-Se: Dunn, P=0.046
Pf (n=5)
Rr (n=1)
Se (n=3)
Sr elemental mass fraction in otoliths (µg g-1)
Species

Figure A.4: Comparison of Sr elemental mass fractions of different fish species from the same habitats, with results of Kruskal-Wallis and pairwise Dunn post-hoc Tests or Mann-Whitney *U*-Tests (Aa=*Anguilla anguilla*, Ab=*Abramis brama*, Bb=*Barbus barbus*, Cspp=*Coregonus spp* El=*Esox lucius*, Pf=*Perca fluviatilis*, Lle=*Leucisus leuciscus*, Ll= *Lota lota*, Rr=*Rutilus rutilus*, , Sc=*Squalius cephalus*, Se=*Scardinius erythrophthalmus*, St=*Salmo trutta*, Tt=*Thymallus thymallus,* Titi=*Tinca tinca*). Comparisons with only one individual of St in Lake Chiemsee were not considered in post hoc test.
